# Altered Frontoparietal, Temporal and Sensorimotor Structure-Function Coupling and Its Genetic Underpinnings in Bipolar Disorder

**DOI:** 10.64898/2025.12.18.695223

**Authors:** Shir Dahan, Lara Ressin, Vilte Baltramonaityte, Emma Corley, Aodán Laighneach, Esther Walton, Pilib Ó Broin, Dara M Cannon

## Abstract

**Background:** Structure-function coupling quantifies how strongly structural connectivity supports functional communication across brain regions. Investigating structure-function coupling in bipolar disorder, a condition marked by dysconnectivity, may elucidate underlying neural mechanisms. We examined regional structure-function coupling in bipolar disorder and its genetic underpinnings to characterize network-level disruptions.

**Methods:** Regional structure-function coupling was estimated in UK Biobank participants using edge-wise regression between measures of structural connectivity and functional connectivity. Bipolar disorder (n=163) and controls (n=326) were age and sex-matched and compared using general linear models. To explore the genetic basis of these structure-function coupling alterations, genome-wide association studies (GWAS) were conducted in an independent UK Biobank sample (n=38,190) for regions showing significant group differences.

**Results:** Structure-function coupling demonstrated a unimodal-to-transmodal gradient with highest coupling evident in visual regions (R^2^=0.29) and lowest in the insula (R^2^=0.03). The Bipolar disorder group showed higher structure-function coupling in the temporal pole and superior frontal gyrus (β=0.269; β=0.211) and lower coupling in the supramarginal, precentral, and postcentral gyri and the frontal pole (β=-0.206 to -0.275). GWAS identified seven significant loci, with mapped genes (e.g., *KAT6B*, *INPP5A*, *PLCE1*) involved in neuronal development and cellular signaling.

**Conclusion:** Bipolar disorder showed altered structure-function coupling across regions implicating multiple networks. These alterations suggest stronger structure-function alignment in limbic and attention networks and weaker alignment in sensorimotor, executive, frontoparietal, and salience regions relative to controls, potentially disrupting flexible polysynaptic communication. Altered coupling may relate to genetic variation affecting neurodevelopment, neuronal signaling, and synaptic plasticity. Together, these findings offer novel insight into the architectural contributions to the pathophysiology underlying bipolar disorder.

## 1 Introduction

Bipolar disorder (BD) is a neuropsychiatric disorder characterized by recurring episodes of mania or hypomania and depression,^1^ affecting 1-3% of the global population.^2^ Dysregulation of emotion and cognition, supported by abnormalities in executive control and limbic cortices, is a core feature of BD.^3^ Neural dysconnectivity is known to underlie the disorder, with reported alterations in both structural and functional connectivity.^4, 5^ Structure-function coupling (SFC), which quantifies how white-matter connectivity relates to correlated functional activity of grey matter regions, offers a framework for understanding the disrupted network organization characteristic of BD. Here, we assess whether BD is associated with regional SFC. As genetic variation has been related to both BD^6^ and SFC^7^ we further investigate the genetic contribution to SFC changes associated with BD.

SFC reflects the degree of alignment between structural connectivity (SC) and functional connectivity (FC),^8^ indicating how anatomical pathways support or adapt to functional interactions.^9, 10^ SFC can be estimated using different approaches, including statistical and biophysical methods^11^ and at different scales (global, meso-scale, local/nodal).^12^ Nodes can be defined, for example, from atlas parcellations^13^ or independent component analysis (ICA).^14^ Differences in SFC may reflect plasticity, myelination, and synaptic pruning related to developmental trajectories that influence healthy and neuropsychiatric phenotypes^8^ as well as shorter-term adaptive processes.^15^ SFC has been implicated in cognitive function^16^ and multiple psychiatric disorders,^12^ including major depressive disorder,^17^ schizophrenia,^18^ and BD.^14^

Mapping structural connections can provide measures related to integration, segregation, and influence, capturing properties of network topology, communication, and organization.^19^ These features can be examined as predictors of FC and show better predictive value in regional compared to global models.^20^ Regional SFC is theorized to vary on a hierarchical gradient,^13^ with higher coupling in evolutionarily more conserved unimodal cortical regions of specific specializations, including sensorimotor and visual regions, and lower coupling in association and poly-sensory transmodal regions, including the default mode network (DMN) and limbic regions.^7, 13, 20^ This gradient is attributed to the evolutionary expansion of association cortices, which decouple neural signaling from structural constraints^13, 21, 22^ and are characterized by heightened experience-dependent plasticity.^23^

Structural and functional graphs differ in spatial and temporal properties. Structural connectomes are estimated using tractography of diffusion-weighted MRI (DWI), with edges representing the strength or existence of connections between parcellated grey matter nodes defined using T1-weighted MRI. They are sparse, temporally stable, and topologically optimized for cost-efficiency, dominated by short-range projections.^24^ Functional graphs derived using functional MRI (fMRI) reflect statistical dependencies between regional blood-oxygen-level-dependent signal time series^25^ and tend to be dense and temporally dynamic.^19^ While FC partially reflects an underlying neuroanatomical backbone, many neural interactions arise from indirect polysynaptic pathways.^10^ Consequently, structure-function correspondence is partial.^26^

Both SC and FC are altered in BD^4, 5, 27^ with evidence for changes in several resting-state networks (RSNs), including the DMN, limbic, ventral attention, frontoparietal, and thalamus networks.^28, 29^ Compared to controls, BD show lower global efficiency^30, 31^ and longer characteristic path length^32^ in structural networks, and higher global efficiency and shorter characteristic path length in functional networks.^33^ These trends suggest decoupling or reorganization of structure-function relationships in BD. These findings, together with the regional variability observed in SFC, highlight the value of multimodal approaches for capturing the complexity of connectivity alterations in BD.

Global SFC findings in BD are inconsistent,^31, 34, 35^ likely reflecting regional variability that is obscured by single global estimates.^13^ At a meso-level, reflecting subnetwork organization, intra-hemispheric SFC was lower in euthymic BD compared to controls,^35^ long-range connection SFC was increased in BD offspring,^36^ and dynamic SFC from ICA was altered in BD participants with varying symptom severity.^14^ Locally, the same dynamic SFC study found higher SFC in the middle frontal gyrus and lower SFC in the supramarginal gyrus in BD,^14^ implicating frontoparietal hubs. However, as these findings were based on group-level ICA nodes, representing distributed networks,^37^ subject-specific and anatomically localized effects may have been obscured. Here, we employ an atlas-based parcellation applied at the single-subject level^12^ with a higher node count to better capture individual and regional effects. Given evidence that SFC outperformed SC and FC in predicting BD and other disorders,^12, 35^ examining its alterations may help identify disorder-specific neuroanatomical patterns with translational relevance.

The genetic basis of structure-function interactions remains underexplored. Genome-wide association studies (GWASs) of SC^38^ and FC^39, 40^ found that both modalities are moderately heritable at certain regions, yet their genetic architectures have little to no genetic overlap.^41^ Kinship analysis revealed that regional SFC is heritable, mostly in subcortical, cerebellar, brainstem, and visual regions.^7^ Additionally, a GWAS on SFC, calculated from surface-based connectivity integration, reported heritability in visual, dorsal attention, language, and sensorimotor networks. Notably, SFC of several networks had a genetic correlation with psychiatric and cognitive traits.^42^ Both the kinship and Single-nucleotide polymorphism (SNP)-based heritability studies found SFC heritability to differ from that of SC and FC.^7, 42^ As SFC reflects a relationship between SC and FC and is shaped by neuronal signaling, development, and plasticity,^8^ it is plausibly influenced by distinct gene sets. Considering that BD has a genetic component,^6, 43^ investigating the genetic underpinnings of BD-associated brain phenotypes may clarify how genetic risk translates to connectivity dysfunction.

In this study, we first aimed to characterize regional SFC across the whole brain and identify regions that are altered in individuals with BD compared to controls, using data from the UK Biobank (UKB). We adopted a data-driven approach, assessing SFC across all brain regions. We then aimed to examine the genetic contribution to regional SFC in regions showing significant diagnosis-group differences, performing a GWAS in an independent UKB sample. We hypothesized SFC alterations in hubs previously implicated in core impairments of BD, and although BD-related regions are often transmodal and therefore less heritable, we hypothesized that genetic variants associated with neural development may underlie altered SFC in BD relative to controls.

## 2 Methods

### 2.1 Study Design

The UKB project is a large-scale prospective cohort study conducted across the UK. The UKB data includes genetic, phenotypic, and health-related information for more than half a million participants. Neuroimaging data have been collected from 2014 and are provided for ∼40,000 participants. The studies adhered to the Declaration of Helsinki. Written informed consent was obtained from all participants, and the North West Multicenter Research Ethics Committee approved this study (No. 11/NW/0382).

### 2.2 Participants

BD inclusion criteria were defined as having an ICD-10 diagnosis of bipolar affective disorder (Chapter V, Code F31, Data-Field 41270) or fulfilling the UKB criteria for BD-I or II diagnosis (Data-Field 20122) or self-reported BD (Data-Field 20002). Participants with any neurological disorder (Chapter VI, Data-Field 41270) were excluded. Control participants were selected from the remaining UKB cohort and age- and sex-matched to the BD group at a 2:1 ratio, excluding individuals with psychiatric or neurological diagnoses defined by ICD-10 Chapters V or VI (Data-Field 41270).

### 2.3 Image Preprocessing and Analysis

Neuroimaging data acquisition, preprocessing, and quality control were performed and made available by the UKB imaging team. A detailed description of the protocol is available at https://biobank.ctsu.ox.ac.uk/crystal/crystal/docs/brain_mri.pdf and elsewhere.^44, 45^ This study used the UKB release of structural connectomes and regional functional time series (Category 509) generated and described by Mansour *et al.* (2023).^46^ For full details, see Supplementary Materials: *‘Data Acquisition and Preprocessing’* and *‘Structural Connectivity Matrices’*. Cortical parcellation was performed using the Schaefer 7-network atlas with 200 or 500 regions,^47^ combined with the Melbourne subcortical atlas comprising 16 or 54 regions,^48^ respectively, and mapped to each participant’s native space. SC matrices were weighted using spherical-deconvolution informed filtering of tractograms (SIFT2).^49^ FC matrices were constructed from standardized regional time series, using Pearson and partial correlation, for each pair of regions (Python 3.9, Nilearn 0.10.4). Pearson correlation captures total statistical dependence, including shared variance mediated by other regions, whereas partial correlation estimates direct conditional dependencies by regressing out the influence of all other nodes. Using both measures allowed assessment of SFC across complementary representations.

### 2.4 Regional Structure-Function Coupling

We quantified SFC (Figure 1) for each participant and each region using a multilinear regression with the predicted variable as the vector of FC edges from each region *i* to all other regions *j* ≠ *i*, and the predictors as standardized SC vectors of structural properties from the same region *i* to all others *j* ≠ *i*.^13^ SC predictors included shortest path length, communicability, and mean first-passage time (MFPT), all weighted, with Euclidean distance added as an additional, geometry-based predictor (Supplementary Materials: *‘Structural Predictors of Functional Connectivity’*). Weighted communicability was used due to the sparse nature of structural connectomes, along with MFPT, and Euclidean distance, which were found to explain more variance in FC compared to other topological predictors.^20^ We then calculated the adjusted R^2^ between the predicted and observed FC vectors as an estimate of the regional SFC, resulting in 216 and 554 adjusted R^2^ values per participant, for FC from both Pearson and partial correlations (Nilearn 0.10.4)

**Figure 1.**
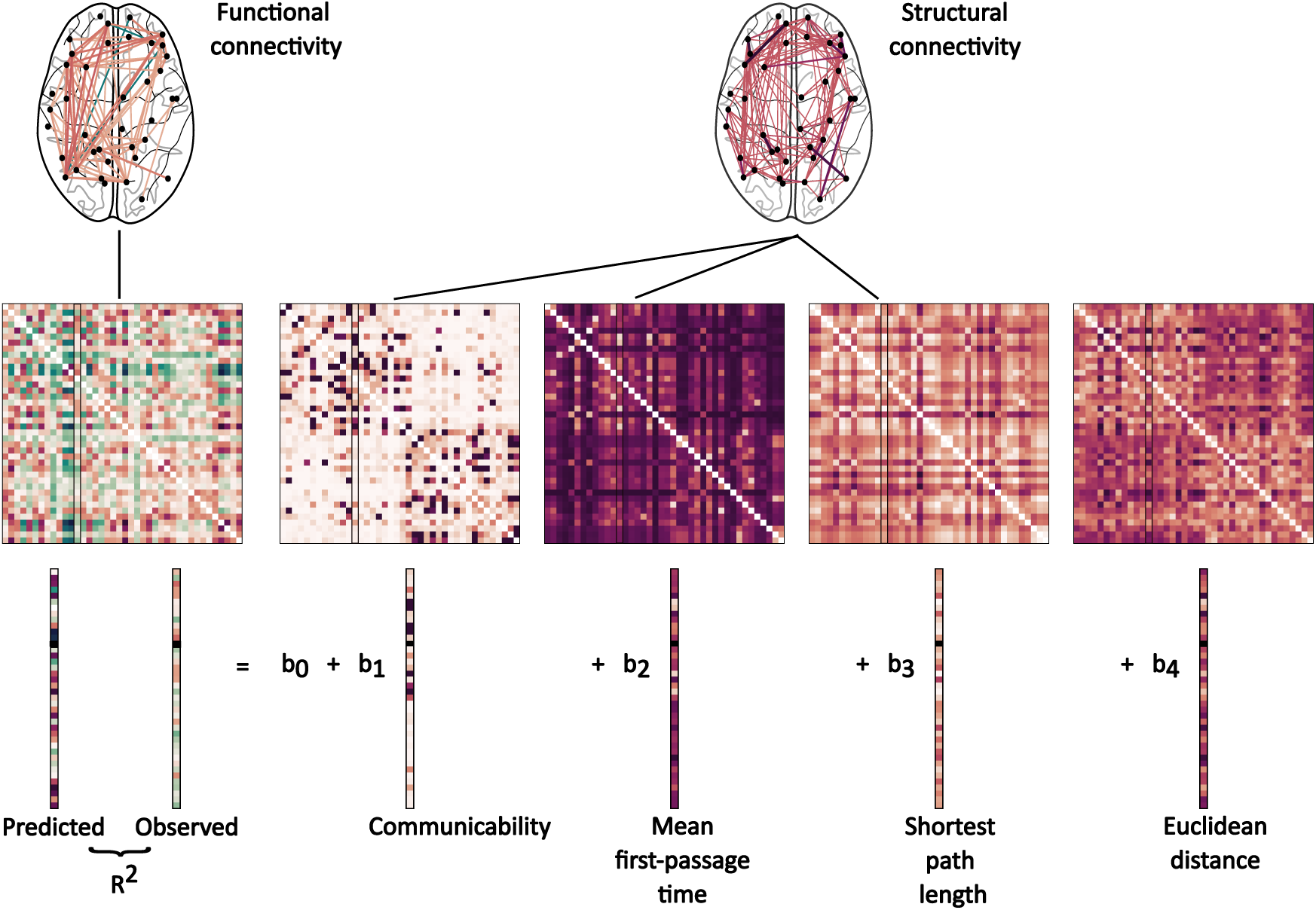
Estimating regional structural connectivity to functional connectivity coupling Figure 1: Structure-function coupling was calculated for each individual and region using multilinear regression, expressed as the goodness of fit (adjusted R^2^) for the prediction of functional connectivity (Pearson/partial) from structural connectivity. Predictors derived from the SIFT2-weighted structural connectome included Communicability, Mean First-Passage Time, and Shortest Path Length, with Euclidean distance added as an additional geometry-based predictor.

### 2.5 Within and Between Network Analysis

We quantified SFC at within-network and between-network levels. Cortical regions were assigned to RSNs according to the Schaefer 7-network atlas, and all subcortical regions from the Melbourne atlas were grouped. Each network is labeled *N_k_*, where *k* = 1*, …, K* indexes the different networks. For each region *i* ∈ *N_k_*, within-network analyses included edges connecting to all other regions *j* ∈ *N_k_, j* ≠ *i* whereas between-network analyses included edges connecting to all regions *j* ∈*/ N_k_*. SFC was then calculated separately for within and between-network connections.^7^

### 2.6 Resting-State Network Analysis

To examine SFC at a network level,^7^ we averaged all regional SFC values assigned to the same RSN in the Schaefer 7-network atlas and all subcortical regions from the Melbourne atlas.

### 2.7 Nodal Degree and Strength

As a supplementary analysis to estimate the proportion of SFC variability attributable to direct local connectivity, we tested the relationship of regional SFC with its degree/strength.^13^ For each region, structural degree was calculated as the number of existing structural connections from that region, excluding self-connections. Structural and functional strength were calculated as the sum of structural and absolute functional connection weights, respectively, excluding self-connections.

### 2.8 Statistical Analyses

Group differences in SFC between BD and controls were assessed using general linear models with a gamma distribution and log link, with diagnosis-group as the predictor of interest and regional SFC as the outcome. Models adjusted for age, sex, intracranial volume, mean resting-state fMRI motion, mean DWI motion, and imaging site. Results were FDR-corrected for the number of regions (α=0.05). This was performed across both parcellation resolutions and FC correlation methods. Associations between SFC and structural degree, structural strength, and functional strength were examined using linear regression (statsmodels 0.14.5).

### 2.9 Genome-Wide Association Analyses

GWASs were performed in an independent UKB sample (previous cohort excluded) to assess the association of genetic variants with regional SFC (R^2^ values) that demonstrated consistent effect on BD associations across both parcellation scales, using REGENIE (v4.1).^50^ REGINIE uses a mixed-model framework that accounts for population structure and kinship. Therefore, we retained related participants across all ancestries to maximize statistical power. Step 1 was run using genotyped array data (Data-Field 22418). A whole-genome model was fitted using ridge regression to estimate the genetic component of each SFC trait. The resulting prediction model was used to compute polygenic predictions using a leave-one-chromosome-out (LOCO) approach. Step 2 was performed using imputed genotype data (Data-Field 21008) with linear regression between variants and the phenotype while incorporating Step 1 LOCO predictions. Covariates in both steps included sex, age, age squared, imaging site, mean resting state fMRI head motion, mean absolute head motion from DWI, total intracranial volume, and the first 20 genetic principal components to account for population structure and ancestry differences. GWAS follow-up analyses were performed with FUMA v1.5.2.^51^ For quality control procedures and imputation steps, see Supplementary Materials: *‘Genome-Wide Association Studies and Post Analyses’*.

### 2.10 Heritability and Genetic correlations

We conducted *post-hoc* exploratory Linkage disequilibrium (LD) Score Regression (LDSC v1.0.1)^52, 53^ to estimate SNP-based heritability for SFC traits and to assess their genetic correlation with a set of 32 relevant GWASs, including MRI-derived SC and FC of RSNs, psychiatric disorders, including BD, and cognitive traits (Table S1). Only publicly available summary statistics with established quality control and appropriate sample sizes of European ancestry were included. LD scores were calculated based on the 1000 Genomes Project European reference panel.^54^

## 3 Results

### 3.1 Participants

The BD (n=163) and control (n=326) groups were matched for age and sex (2:1 ratio). BD participants had lower high school level educational attainment compared to controls (Table 1).

**Table 1:**
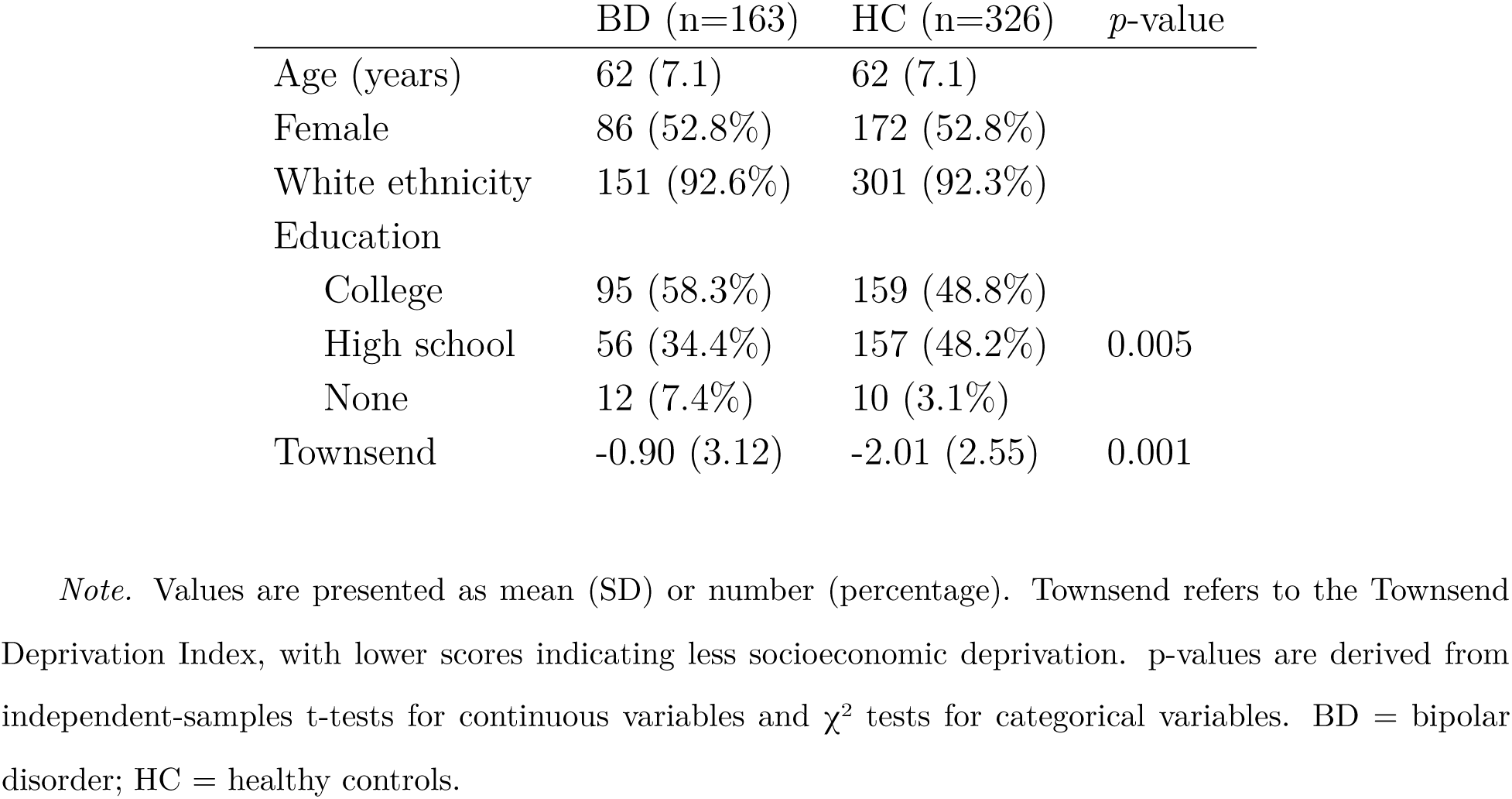
Demographics of the group-comparison cohort

### 3.2 Regional Structure-Function Coupling

SFC across the brain (Figure 2) showed an organized, symmetrical gradient in each hemisphere. SFC was highest in primary and higher-order visual and sensorimotor regions (Table 2) and lowest in salience and limbic hubs and subcortical regions (Table 3). Between-network regional SFC was predominantly high in parietal and frontal areas and lowest in limbic and visual regions (Figures S1, Tables S2-S3). Within-network regional SFC was higher than both overall and between-network regional SFC and did not exhibit a pronounced spatial organization, despite variability in coupling strength (Figures S2, Tables S4-S5). SFC values within regions exhibited a positive skew thereby supporting the use of a gamma log-linked GLM to compare between diagnosis groups. Associations with structural degree and structural strength were small, whereas functional strength showed consistently stronger positive associations with SFC (Figures S3-S4, Table S6).

**Figure 2.**
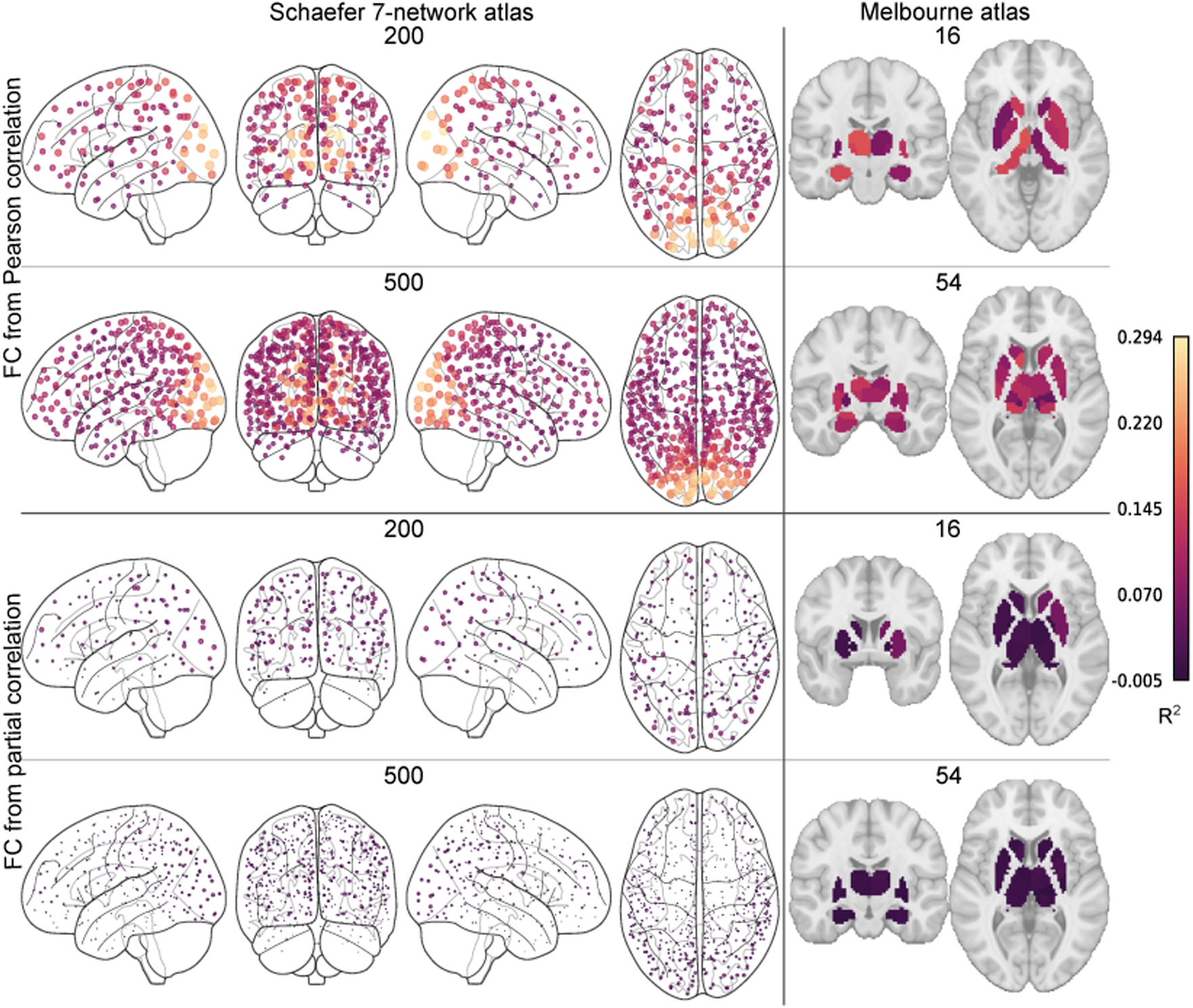
Regional structure-function coupling across cortical and subcortical regions Figure 2: Structure-function coupling (R^2^) was computed in the Schaefer 7-network atlas and subcortical regions in the Melbourne atlas for 200 or 500 cortical and 16 or 54 subcortical regions, with FC estimated using Pearson or partial correlations. Larger and lighter dots indicate stronger coupling. FC = functional connectivity.

**Table 2:**
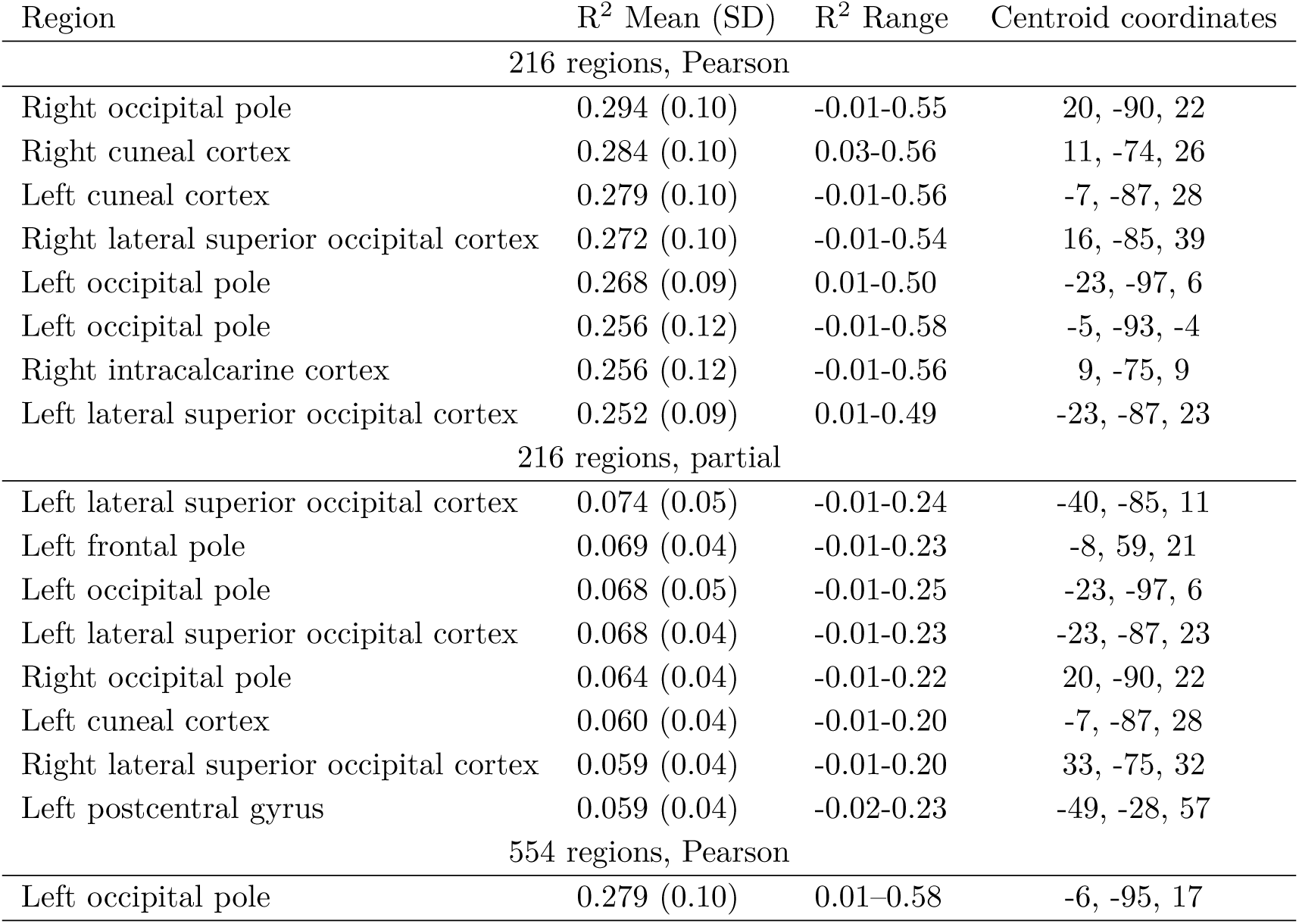
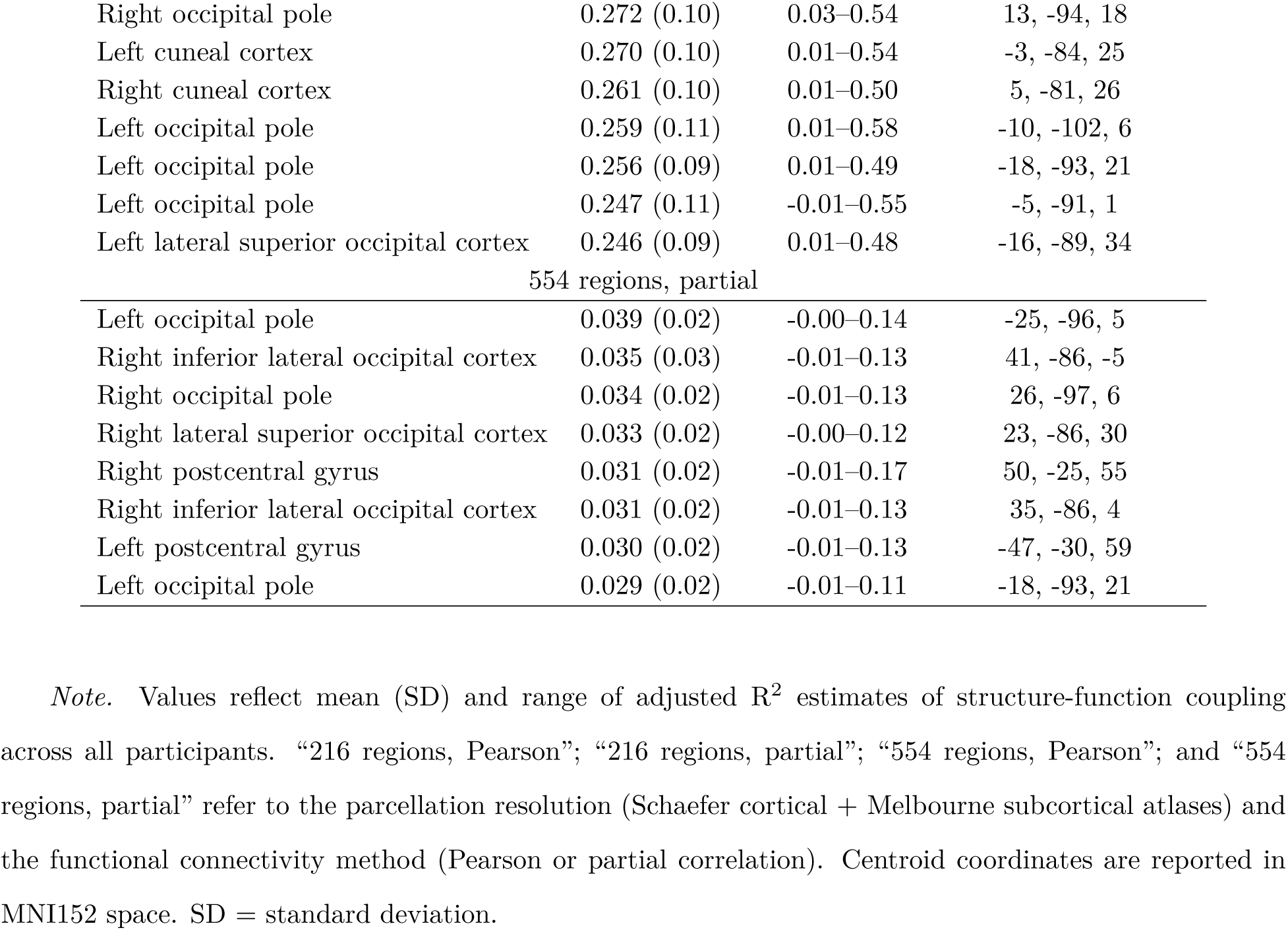
Regions with the highest SC-FC coupling

**Table 3:** Regions with the lowest SC-FC coupling

| Region | $R^2$ Mean (SD) | $R^2$ Range | Centroid coordinates |
| --- | --- | --- | --- |
| 216 regions, Pearson |  |  |  |
| Right hippocampus | 0.033 (0.04) | -0.02-0.22 | 28, -22, -14 |
| Left insular cortex | 0.035 (0.04) | -0.02-0.40 | -33, 20, 5 |
| Right anterior thalamus | 0.036 (0.04) | -0.02-0.23 | 10, -14, 8 |
| Right insular cortex | 0.037 (0.04) | -0.02-0.30 | 41, 6, -15 |
| Right posterior thalamus | 0.038 (0.04) | -0.02-0.22 | 16, -26, 2 |
| Left posterior thalamus | 0.040 (0.04) | -0.02-0.22 | -14, -26, 2 |
| Left hippocampus | 0.040 (0.04) | -0.02-0.19 | -26, -22, -14 |
| Right posterior middle temporal gyrus | 0.041 (0.03) | -0.01-0.19 | 61, -13, -21 |
| 216 regions, partial |  |  |  |
| Left globus pallidus | -0.005 (0.01) | -0.02-0.04 | -18, -4, -2 |
| Left nucleus accumbens | -0.004 (0.01) | -0.02-0.05 | -10, 14, -6 |
| Right nucleus accumbens | -0.004 (0.01) | -0.02-0.04 | 12, 14, -6 |
| Right globus pallidus | -0.004 (0.01) | -0.02-0.07 | 20, -4, -2 |
| Right anterior cingulate gyrus | -0.004 (0.01) | -0.02-0.07 | 5, 3, 30 |

Table 3: Regions with the lowest SC-FC coupling (Continued)
|  |  |  |  |
| --- | --- | --- | --- |
| Right posterior thalamus | -0.003 (0.01) | -0.02-0.05 | 16, -26, 2 |
| Right amygdala | -0.003 (0.01) | -0.02-0.04 | 24, -4, -18 |
| Left anterior cingulate gyrus | -0.003 (0.02) | -0.02-0.31 | -3, 4, 30 |
| 554 regions, Pearson |  |  |  |
| Right medial dorsoanterior thalamus | 0.028 (0.03) | -0.01-0.13 | 10, -24, 12 |
| Left medial dorsoanterior thalamus | 0.032 (0.03) | -0.01-0.17 | -8, -24, 12 |
| Right anterior cingulate gyrus | 0.034 (0.03) | -0.01-0.22 | 7, 8, 41 |
| Right hippocampus body | 0.034 (0.03) | -0.01-0.19 | 30, -28, -10 |
| Right lateral dorsoanterior thalamus | 0.035 (0.03) | -0.01-0.15 | 16, -20, 12 |
| Right medial ventroposterior thalamus | 0.036 (0.03) | -0.01-0.23 | 10, -24, 2 |
| Right anterior cingulate gyrus | 0.038 (0.03) | -0.01-0.21 | 6, -12, 30 |
| Left posterior cingulate gyrus | 0.038 (0.03) | -0.01-0.26 | -5, -23, 29 |
| 554 regions, partial |  |  |  |
| Left anterior globus pallidus | -0.002 (0.0) | -0.01-0.03 | -16, 0, -2 |
| Left posterior globus pallidus | -0.002 (0.0) | -0.01-0.02 | -22, -8, -2 |
| Right subcallosal cortex | -0.002 (0.0) | -0.01-0.02 | 4, 24, -22 |
| Right posterior temporal fusiform cortex | -0.002 (0.0) | -0.01-0.02 | 40, -15, -31 |
| Right posterior inferior temporal gyrus | -0.002 (0.0) | -0.01-0.01 | 45, -31, -23 |
| Left posterior temporal fusiform cortex | -0.002 (0.0) | -0.01-0.02 | -32, -24, -26 |
| Left posterior temporal fusiform cortex | -0.002 (0.0) | -0.01-0.01 | -39, -16, -31 |
| Left anterior inferior temporal gyrus | -0.002 (0.0) | -0.01-0.02 | -46, -5, -41 |

*Note.* Values reflect mean (SD) and range of adjusted $R^2$ estimates of structure-function coupling across all participants. “216 regions, Pearson”; “216 regions, partial”; “554 regions, Pearson”; and “554 regions, partial” refer to the parcellation resolution (Schaefer cortical + Melbourne subcortical atlases) and the functional connectivity method (Pearson or partial correlation). Centroid coordinates are reported in MNI152 space. SD = standard deviation.

Averaged across regions assigned to the same RSNs, the visual network consistently showed the highest SFC, whereas the limbic network and subcortex exhibited the lowest (Figure 3, Table S7). This pattern also appears in the between-network analysis (Figure S5, Table S8). In the within-network analysis, average SFC values varied across networks but did not exhibit a stable network-level hierarchy across parcellations (Figure S6, Table S9).

**Figure 3.**
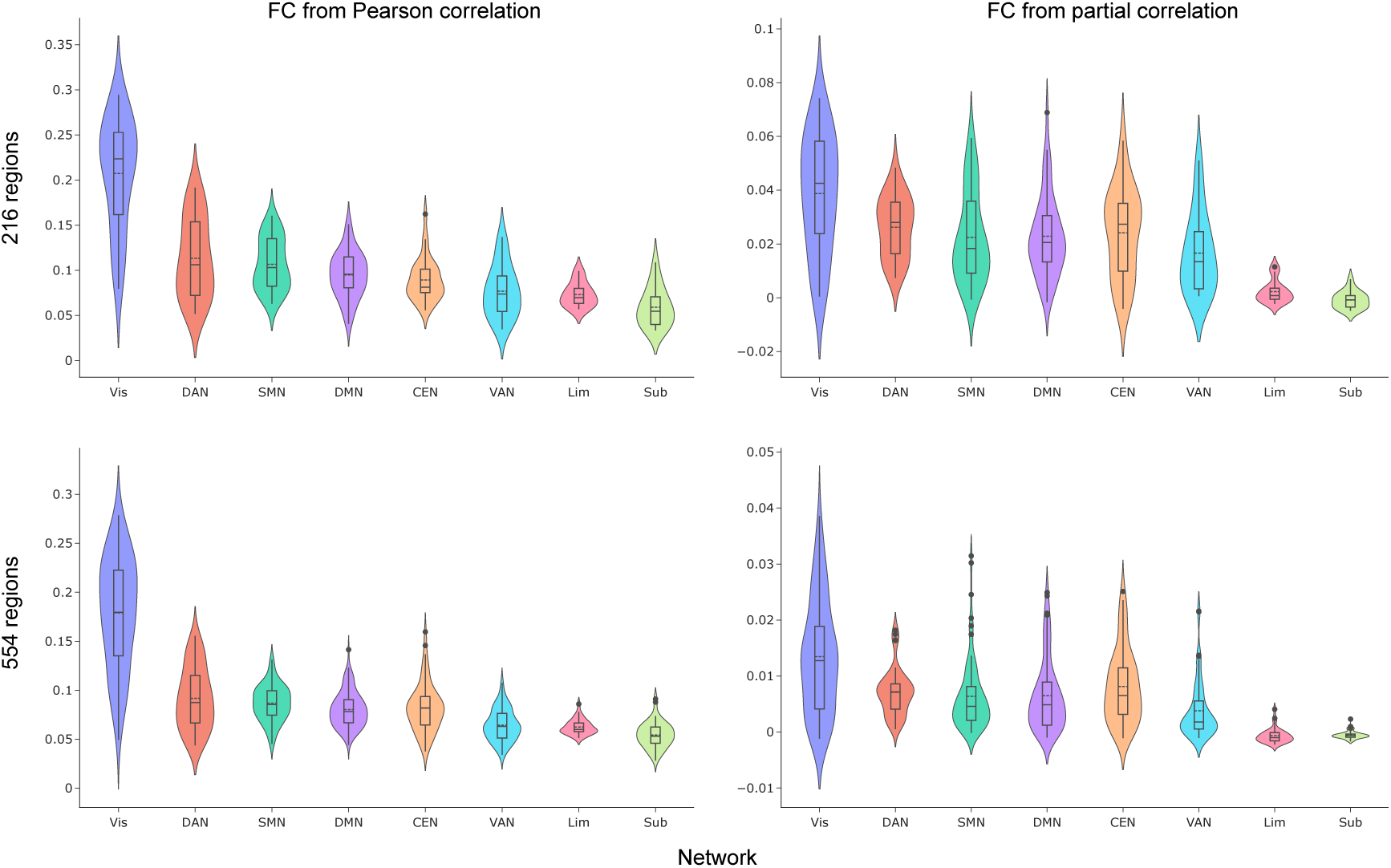
Regional structure-function coupling (R^2^) averaged across resting-state networks Figure 3: Structure-function coupling (R^2^) was averaged across resting-state networks as assigned by the 200 or 500-region Schaefer 7-network atlas, and with 16 or 54 subcortical regions in the Melbourne atlas, with functional connectivity calculated using Pearson or partial correlation. CEN = central executive; DAN = dorsal attention; DMN = default mode; FC = functional connectivity; Lim = limbic; SMN = sensorimotor; Sub = subcortical; VAN = ventral attention; Vis = visual.

### 3.3 Structure-Function Coupling Differences in Bipolar Disorder

Several regions showed significant group differences in SFC (Figure 4, Table 4). Higher coupling in the BD group was observed in the left temporal pole and in the right superior frontal gyrus. Lower coupling in BD was found in the left precentral gyrus, the left anterior supramarginal gyrus, the right frontal pole, and the right postcentral gyrus. Of these regions, only the right postcentral gyrus showed an inconsistent direction of effect between resolutions.

**Figure 4.**
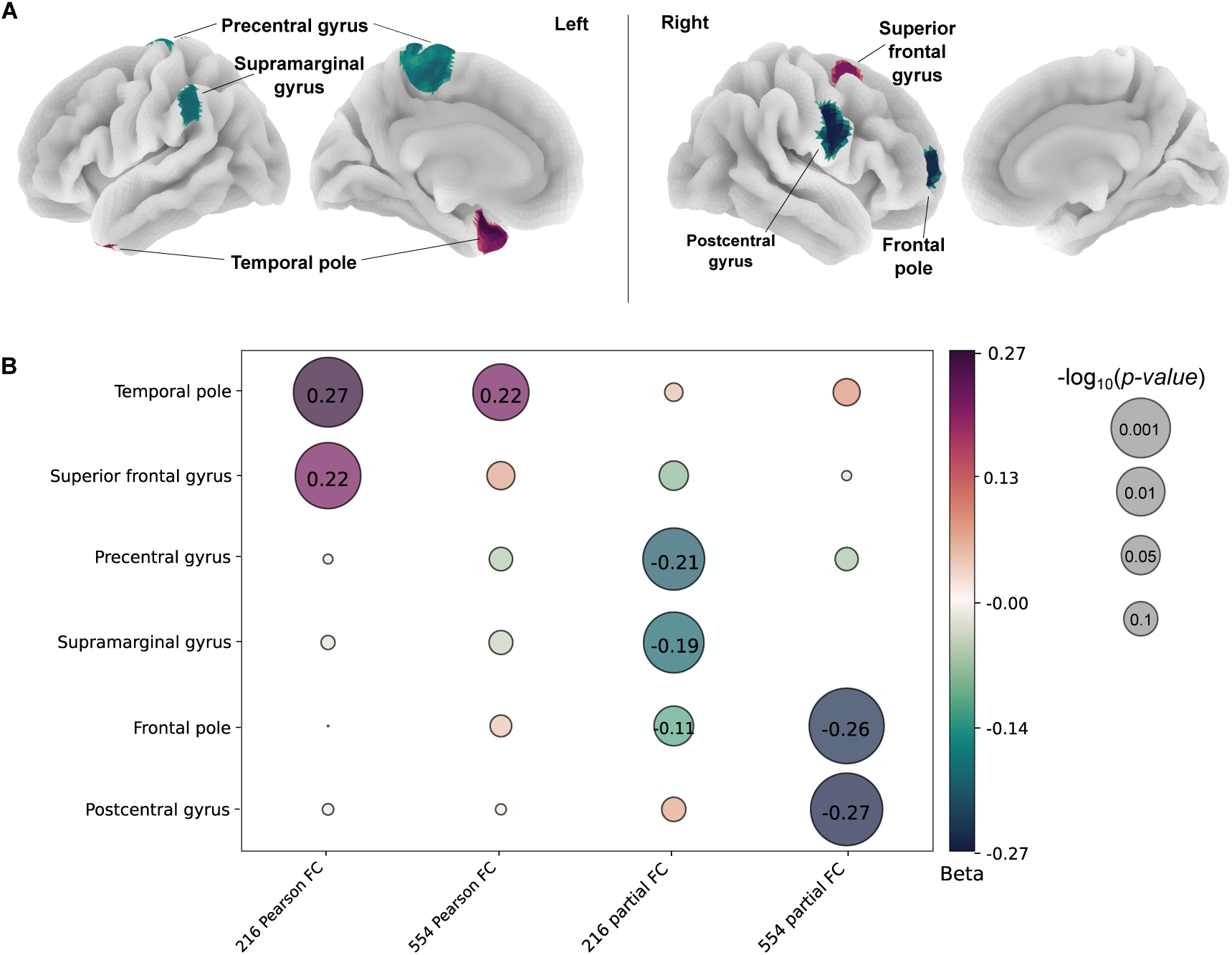
Regional structure-function coupling significant differences between bipolar disorder and control groups Figure 4: Regions with significant difference in structure-function coupling between bipolar disorder and control groups A) across all and B) compared between parcellations and functional connectivity types. The color represents the beta, and the bubble size represents the − log_10_(*p*) values with the raw *p*-value annotated inside the bubble. FC = functional connectivity.

**Table 4:** Regional structure-function coupling differences between bipolar disorder and control groups

| Region | N regions | Centroid coordinates | Direction | $\beta$ | $p$ | $p_{FDR}$ | 95% CI |
| --- | --- | --- | --- | --- | --- | --- | --- |
| FC from Pearson correlation |  |  |  |  |  |  |  |
| Left temporal pole [Lim] | 216 | -28, 10, -34 | BD>HC | 0.267 | 0.001 | 0.018* | 0.13, 0.40 |
|  | 554 | -27, 10, -33 | BD>HC | 0.236 | 0.001 | 0.305 | 0.10, 0.37 |

Table 4: Regional structure-function coupling differences between bipolar disorder and control groups (Continued)
|  |  |  |  |  |  |  |  |
| --- | --- | --- | --- | --- | --- | --- | --- |
| Right superior frontal | 216 | 26, 7, 58 | BD>HC | 0.211 | 0.001 | 0.026* | 0.10, 0.32 |
| gyrus [DAN] | 554 | 25, 9, 52 | BD>HC | 0.07 | 0.210 | 0.835 | 0.08, 0.19 |
| FC from partial correlation |  |  |  |  |  |  |  |
| Left precentral gyrus [SMN] | 216 | -5, -29, 67 | BD<HC | -0.227 | 0.001 | 0.029* | -0.34, -0.11 |
|  | 554 | -4, -26, 58 | BD<HC | -0.064 | 0.340 | 0.839 | -0.19, 0.067 |
| Left anterior supramarginal | 216 | -60, -39, 36 | BD<HC | -0.206 | 0.001 | 0.029* | -0.30, -0.10 |
| gyrus [VAN] | 554 | -62, -36, 36 | BD<HC | -0.004 | 0.953 | 0.993 | -0.14, 0.13 |
| Right frontal pole [CEN] | 216 | 29, 58, 5 | BD<HC | -0.153 | 0.006 | 0.187 | -0.26, -0.04 |
|  | 554 | 30, 57, 6 | BD<HC | -0.275 | 0.001 | 0.002* | -0.39, -0.16 |
| Right postcentral | 216 | 58, -5, 31 | BD>HC | 0.070 | 0.306 | 0.764 | -0.06, 0.20 |
| gyrus [SMN] | 554 | 64, -11, 29 | BD<HC | -0.274 | 0.001 | 0.007* | -0.40, -0.15 |

*Note.* Results of comparisons in regional structure-function coupling across both parcellation resolutions. Each region is presented with centroid MNI-152 coordinates, direction of group difference, effect estimate ( $\beta$ ), $p$ -value, FDR-corrected $p$ -value, and confidence interval. Findings surviving FDR correction are marked with an asterisk. Rows show the same anatomical region across the 216 and 554-region parcellations for comparison. Network assignment (Schaefer atlas) shown in brackets []. CEN = central executive; DAN = dorsal attention; FC = functional connectivity; Lim = limbic; SMN = sensorimotor; VAN = ventral attention.

### 3.4 Genome-Wide Association Studies

Genome-wide association analyses in an independent cohort (n=38,190; mean age 55±8 years; 53% female; 87% Caucasian; 76% unrelated) were performed on SFC in five regions showing consistent SFC difference directionality across parcellations: the left temporal pole, precentral gyrus, supramarginal gyrus, and the right superior frontal gyrus and frontal pole. Genome-wide significant associations (*p <* 5 × 10*^−^*^8^) were identified for SFC in three regions: precentral gyrus, frontal pole, and temporal pole, with no significant loci detected for the supramarginal or superior frontal gyri. Five independent loci were associated with left precentral gyrus SFC, located on chromosomes 2, 5, 8, 10, and 11. One locus on chromosome 10 was associated with temporal pole SFC, and one locus on chromosome 10 was associated with frontal pole SFC (Figure 5, Tables S10–S11; Supplementary Materials: *’Genome-Wide Association Studies’*).

**Figure 5.**
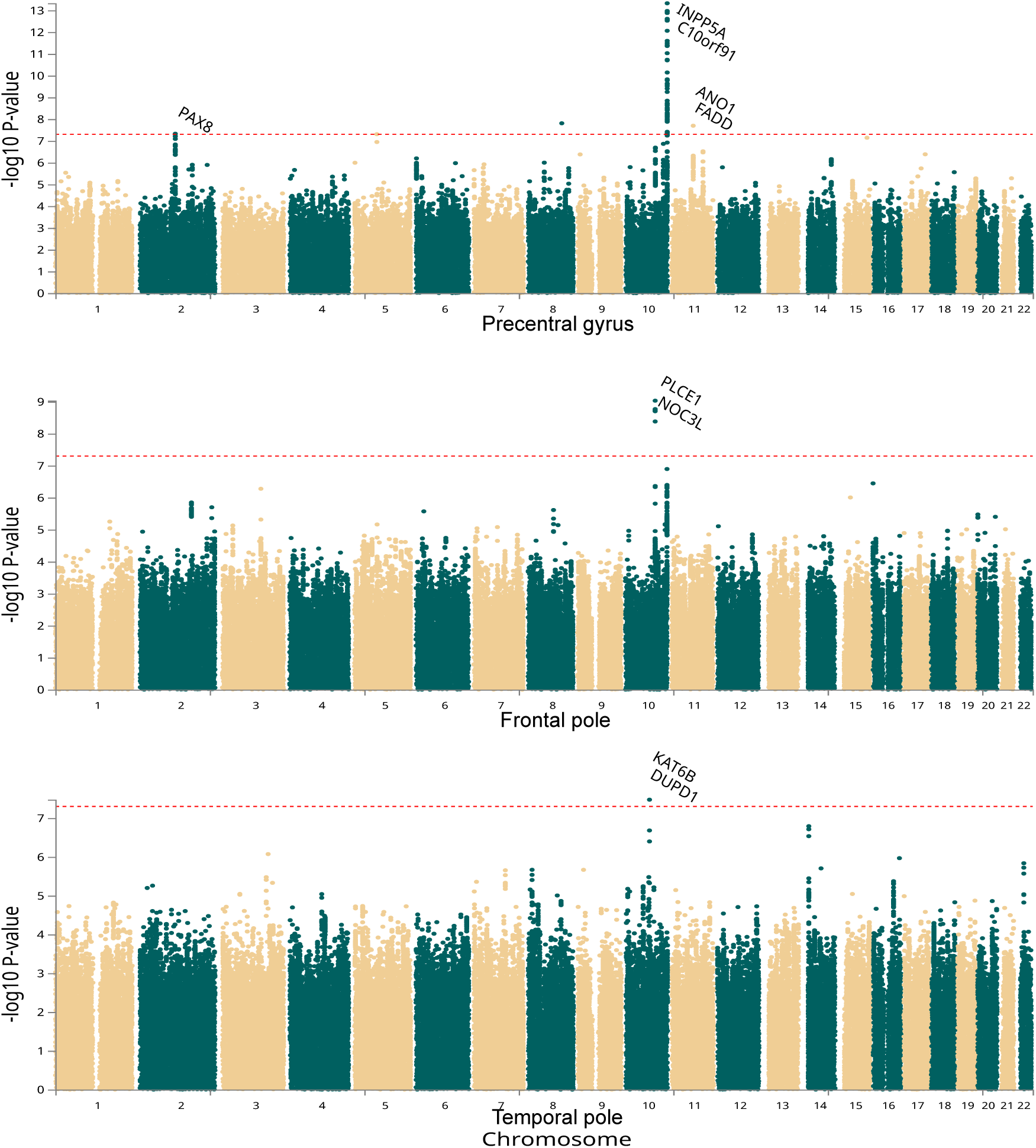
Manhattan plot of genome-wide association results for structure-function coupling regions associated with bipolar disorder Figure 5: Manhattan plots are shown for the regions that had significant single-nucleotide polymorphisms: precentral gyrus, frontal pole, and temporal pole. Each plot displays SNP-level − log_10_(*p*) values mapped to genomic position. The red dotted line denotes the genome-wide significance threshold (*p <* 5×10*^−^*^8^). Significant loci, where present, correspond to variation associated with regional structure-function coupling. Genes mapped by FUMA SNP2GENE with positional or eQTL mapping are annotated.

Expression quantitative trait locus (eQTL) analyses (PsychENCODE and GTEx v8) showed that across brain tissues, precentral gyrus SFC risk alleles were associated with increased *INPP5A* and decreased *PAX8* and *PSD4* expression, while frontal pole SFC risk alleles were associated with increased *NOC3L* and *PLCE1* expression; no significant eQTLs were identified for temporal pole SFC. Functional mapping using FUMA revealed that several lead SNPs from the precentral gyrus GWAS had prior associations in the GWAS Catalog, including *rs7914305* (*INPP5A*) and *rs12762160* (*C10orf91*) previously linked to resting-state fMRI traits. *rs2863957* (*PAX8*) has been associated with sleep duration, insomnia-related depressive symptoms, and brain functional connectivity, while *rs5792512* (*ANO1*, *FADD*) has been reported for cardiovascular traits. In the frontal pole, GWAS Catalog overlap included 134 shared associations, predominantly involving neuroimaging, cardiometabolic, blood, and immune traits, whereas no catalog overlap was observed for temporal pole SFC.

### 3.5 Heritability and Genetic Correlation

Only three regions exhibited sufficient genetic signal to allow heritability estimation: Precentral gyrus SFC showed modest heritability, whereas supramarginal gyrus SFC showed only a small heritable component with some indication of inflation, and frontal pole SFC showed minimal evidence of heritability (Table 5). Exploratory genetic correlation analyses revealed nominally significant associations (*p <* 0.05) only for SFC in the precentral gyrus with several MRI-derived structural and functional connectivity measures, reaction time, feelings of guilt, and age of first depressive episode (Figure S7; Table S1). None survived FDR correction for the number of traits.

**Table 5:** Heritability estimates of genome-wide association analyses

| SFC trait | $h^2$ | SE ($h^2$) | $\lambda_{GC}$ | LDSC intercept ± SE |
| --- | --- | --- | --- | --- |
| Frontal pole | 0.0089 | 0.0125 | 1.01 | $1.0075 \pm 0.0078$ |
| Precentral gyrus | 0.0585 | 0.0137 | 1.06 | $1.0177 \pm 0.0077$ |
| Supramarginal gyrus | 0.0189 | 0.0033 | 1.20 | $1.0927 \pm 0.0301$ |

*Note.* SNP heritability estimates for genome-wide association analyses of regional structure-function coupling. For each trait, the heritability estimate ( $h^2$ ), its standard error, genomic inflation factor ( $\lambda_{GC}$ ), and LDSC intercept with its standard error are shown. Values reflect LDSC results applied to each GWAS. LDSC = Linkage disequilibrium score regression.

## 4 Discussion

This study investigated regional coupling between structural and functional connectivity to extend our understanding of the interplay of neural network configuration and functional dynamics in bipolar disorder. Individuals with BD showed significantly higher SFC in the left temporal pole and right superior frontal gyrus, and lower SFC in the left precentral and supramarginal gyri and the right postcentral gyrus and frontal pole. GWASs revealed genome-wide 7 significant loci for three regions associated with altered SFC in BD.

### 4.1 Regional Structure-Function Coupling Across the Brain

Overall, SFC exhibited a bilaterally symmetric unimodal-to-transmodal gradient, consistent with prior evidence,^13^ reflecting that functional organization follows more closely underlying neuroanatomical connectivity in primary cortices, compared to transmodal regions of diverse functions.^21^ Primary visual regions showed the highest SFC, reflecting tightly organized, predominantly feedforward circuitry supporting early sensory processing.^55^ Limbic, salience, and subcortical regions exhibited the lowest SFC. This pattern is consistent with prior work suggesting that higher-order regions rely on more distributed, long-range connections and exhibit greater functional heterogeneity, leading to weaker structure-function coupling.^56^

The brain network exhibits small-world properties^57^ with densely interconnected and functionally specialized sub-communities, reflecting a balance between local segregation and global integration.^10^ Aligning with this topology, within-network SFC was higher and varied less across networks, consistent with prior reports,^7^ reflecting that dense local clustering supports efficient regional communication, particularly within intrinsic functional networks.^58^ Furthermore, SFC extent varied with parcellation granularity: higher resolution parcellations generally yielded lower SFC values, possibly due to computational scaling or that smaller subregions are less structurally constrained and only converge toward anatomical skeletons when aggregated at coarser scales.

### 4.2 Structure-Function Coupling in Bipolar Disorder

The BD group showed higher SFC in the left temporal pole, a region implicated in limbic, cognitive, and semantic processing^59, 60^ and known to act as a structural and functional integrative hub spanning multiple large-scale networks.^61^ Structural and morphometric alterations of the temporal pole have been reported in BD.^62–65^ Lesions of the left temporal pole have been associated with rapid-cycling BD, as well as mood or psychotic symptoms following lobectomy.^66, 67^ Higher SFC in BD in the superior frontal cortex, another highly interconnected hub,^68^ may relate to dorsal attention network dysfunction.^69^ Reduced superior frontal gray matter volume and altered resting-state activity were reported in BD.^62, 70, 71^ Moreover, greater irritability was linked to reduced cortical thickness, volume, and gyrification of the region.^72^ The region is also implicated in response inhibition and motor urgency, supporting a role in impulsivity regulation.^73^ Increased SFC could indicate more constrained functional dynamics within these integrative regions, potentially contributing to emotion dysregulation.

Lower SFC in the left anterior supramarginal gyrus may reflect altered integration within the ventral attention and semantic circuitry.^74^ Reduced SFC in the precentral and postcentral gyri is consistent with disrupted sensorimotor organization in BD, including reduced gyrification in the precentral gyrus,^75^ weaker within-network connectivity, and altered connectivity with frontoparietal and visual networks.^76, 77^

Sensorimotor circuits are thought to support higher-order cognition during emotional processing. Disruptions of these processes have been proposed to contribute to impaired semantic and emotional integration in affective disorders,^78^ potentially linking the sensorimotor alterations observed here with changes in semantic and limbic regions such as the temporal pole and supramarginal gyrus. Lower SFC in the right frontal pole further implicates large-scale integrative control networks,^79^ consistent with evidence linking atypical synaptic pruning and neurotransmission-related gene expression in this region to BD neurodevelopment.^80^

### 4.3 Genetic Insights into Structure-Function Coupling

The precentral gyrus, a unimodal region, exhibited the strongest heritability, consistent with evidence that unimodal regions are more heritable than transmodal regions.^7, 42^ eQTL effects and chromatin interactions were found in cortical and basal ganglia tissues. These patterns highlight regulatory influences, showing that SFC-associated variants modulate brain transcriptional activity, elevating *INPP5A*, *NOC3L*, and *PLCE1* expression and reducing *PAX8* and *PSD4*. For precentral gyrus SFC, *FADD* has a role in apoptosis, proliferation, and cell cycle progression,^81^ *FOXF2* was previously related to blood-brain barrier maintenance,^82^ and *INPP5A* is involved in cell signaling,^83^ which may affect neuronal development. *PAX8* is implicated in the regulation and production of thyroid hormone^84, 85^ and was also identified in a GWAS of the sensorimotor network FC^41^ of which the precentral gyrus is a part. For the frontal pole, *NOC3L* is involved in the regulation of DNA replication and proliferation,^86^ and *PLCE1* regulates various processes affecting cell growth, differentiation, and gene expression^87^ and was found in a GWAS of limbic network FC.^41^ In the temporal pole, *KAT6B* encodes a histone acetyltransferase involved in chromatin remodeling and neurodevelopment.^88^ In addition, several associated variants overlap with GWAS Catalog entries, predominantly linked to diverse neuroimaging traits.^40, 89–95^ Notably, the mapped genes have not been previously associated with BD.^43^ This may reflect limited power to detect cross-trait overlap, or the exclusion of BD participants from the GWAS, which might have reduced representation of BD-risk alleles. However, given the pleiotropic nature of BD,^96^ the absence of BD-associated genes among SFC loci suggests that regional SFC reflects broad neural processes that may be relevant to the development of BD.

### 4.4 Methodological Considerations

Factors influencing the replicability of SFC findings include SC and FC reconstruction methods, parcellation atlas selection, and the model used to quantify SC-FC correspondence.^12^ We selected SIFT2 weighting for SC due to its strong biological interpretability^49^ and employed parcellation atlases derived from functional gradients, at two parcellation scales, to better align structural and functional data and to assess consistency of findings. In our findings, FC calculated from partial correlation, which measures the direct correlation between two regions while adjusting for all others, yielded lower SFC values compared to Pearson correlation. However, higher SFC values were reported previously using either partial or precision correlation for FC.^25, 97^ These differences may stem from methodological and analytical differences. Additionally, variability has been reported in how SFC maps onto established cortical hierarchies. For example, at an individual level, only weak to moderate associations between SFC and the unimodal-transmodal gradient were found.^98^ Collectively, these considerations highlight the need for careful selection of analysis parameters and standardized, high-quality pipelines to ensure robust and reproducible SFC findings.

### 4.5 Limitations

The linear model used to compute SFC assumed a linear relationship between SC and FC outcome, which may oversimplify the complexity of structure-function relationships. Second, the unequal number of regions across RSNs introduces potential bias when comparing networks and in estimating within and between-network SFC. Third, large within-RSNs variance was observed, such that averaging across regions may obscure important heterogeneity of potentially critical functional significance.

### 4.6 Future Directions

Future work should increase sample sizes to improve discovery power and assess the replicability of group differences and GWAS results. Additionally, examining clinical state-dependent SFC differences could provide biological insight, although such data were unavailable in the UKB cohort. Finally, investigating the potential of SFC as a transdiagnostic marker across neurological and psychiatric disorders could inform specific neurobiological factors involved in the development of BD.

### 4.7 Conclusion

Disrupted communication dynamics of frontoparietal, sensorimotor, limbic, and attentional networks may underlie cognitive and affective symptoms in BD. Structural connections were found to differently constrain functional dynamics in these regions in bipolar disorder, extending our understanding of the neural basis of the disorder. The observed genetic associations underscore the relevance of SFC, reinforcing its value as an integrative phenotype for studying genetics of brain organization. These findings advance understanding of structure-function relationships and large-scale brain network organization in bipolar disorder.

## 5 Statements

### 5.1 Acknowledgments

This research has been conducted using the UK Biobank Resource under Application Number 936468. We sincerely thank the participants for their involvement in this research study. We want to thank Derek Morris and his lab, the Clinical Neuroimaging Laboratory, as well as Mehak Chopra and the Research Ireland Centre for Research Training in Genomics Data Science group. This research was funded by Research Ireland through the Research Ireland Centre for Research Training in Genomics Data Science under Grant number 18/CRT/6214. EW received funding from UK Research and Innovation (UKRI) under the UK government’s Horizon Europe / ERC Frontier Research Guarantee [BrainHealth, grant number EP/Y015037/1] and from Wellcome (reference 315898/Z/24/Z).

### 5.2 Data Availability

While UK Biobank individual-level data cannot be publicly shared, they are accessible to qualified researchers through the UK Biobank Access Management System. GWAS summary statistics generated in this study are available upon reasonable request, and we are glad to share them with interested researchers. Derived, non-identifiable data products can also be shared upon request, in accordance with UK Biobank data-sharing policies.

### 5.3 Author Contribution

Conceptualization SD, EW, POB, DMC; methodology and formal analysis SD, VB, EM, AL, EW, POB, DMC; resources EW, POB, DMC; data curation SD; writing original draft SD, DMC; writing, reviewing, and editing all authors; visualization LR, SD; supervision EW, POB, DMC; funding acquisition POB, DMC.

