## Supplementary Materials for "Altered Frontoparietal, Temporal and Sensorimotor Structure-Function Coupling and Its Genetic Underpinnings in Bipolar Disorder"

### METHODS

#### Data Acquisition and Preprocessing:

Data acquisition and preprocessing performed by Alfaro-Almagro et al. (2020) as described in <https://biobank.ctsu.ox.ac.uk/crystal/crystal/docs/brain_mri.pdf>.

All MRI data were acquired with 3T Siemens Skyra scanners using a standard 32-channel head coil. Four assessment centers were used to collect the data. The T1-weighted structural images were acquired using a 3D-MPRAGE acquisition (1 mm isotropic voxel, 256 mm superior-inferior field of view). Preprocessing steps included gradient distortion correction, skull stripping, linear and nonlinear registration to MNI152 standard space, and defacing (Alfaro-Almagro et al., 2018).

Resting-state BOLD data were acquired with a multi-band gradient echo EPI sequence, with an acquisition time of ∼6 min (490 volumes, 2.4 mm isotropic voxels), TE/TR = 39/735 ms, MB = 8, no in-plane acceleration, flip angle 52◦, conventional fat saturation. Preprocessing steps included the FSL MELODIC pipeline (Beckmann and Smith, 2004) (EPI susceptibility distortion correction, GDC, motion correction with FSL MCFLIRT, grand-mean intensity normalization, and high-pass temporal filtering) followed by an ICA + FIX step to suppress remaining artifact components (Alfaro-Almagro et al., 2018).

Diffusion-weighted MRI (DWI) data were acquired using a multi-band spin echo EPI sequence, with an acquisition time of ∼7 min (100 unique diffusion sensitization directions distributed equally across two shells, b-values: 1000, 2000 s∕mm^2^), and five b=0 volumes, 2 mm isotropic voxels (MB = 3, no in-plane acceleration, TE/TR = 92/3600 ms, partial Fourier 6/8, conventional fat saturation). The data were preprocessed with a pipeline consisting of correction for eddy current and head motion followed by GDC. Additionally, FSL’s dti-fit and the NODDI toolbox were used to generate voxel-wise microstructural parameters (Alfaro-Almagro et al., 2018).

#### Structural Connectivity Matrices:

Connectome construction as performed and described by Mansour et al. (2023):

Preprocessing included skull stripping the DWI data to create a brain mask for T1 and DWI alignment, estimation of tissue response functions for WM, GM, and cerebrospinal fluid CSF, fiber orientation distribution estimation with multi-shell, multi-tissue constrained spherical deconvolution, and intensity normalization with bias field correction. Whole-brain tractography involved tissue segmentation for anatomically constrained tractography (ACT) and masking of the GM-WM interface for streamline seeding. Probabilistic tractography (iFOD2) was performed with 10 million streamlines, and streamlines were filtered based on length constraints and ACT priors. SC matrices were constructed by assigning streamlines to brain regions using parcellation data and computing SC measures. Here, we chose to use Fiber Bundle Capacity (FBC) weights. FBC per streamline was estimated with spherical-deconvolution informed filtering of tractograms (SIFT2) (Mansour L. et al., 2023).

#### Structural Predictors of Functional Connectivity:

Weighted shortest path length, the sum of the shortest route between every pair of nodes, was calculated with the inverse SIFT2 weights using the Floyd Warshall Algorithm (netneurotools 0.2.4). Communicability, which reflects the sum of walks of different lengths between node pairs, was computed by applying the matrix exponential to the SIFT2-weighted connectivity matrix and normalizing the result by its largest eigenvalue, and Mean first-passage time (MFPT) was calculated as the expected number of steps a random walker takes to reach a target node for the first time, using the transition probability matrix derived from SIFT2 weights. To reduce nodal bias, MFPT matrices were row-wise min-max normalized to the [0, 1] range.

#### Genome-Wide Association Studies and Post Analyses:

Preprocessing and quality control: For sample quality controls, we excluded participants with gender not matching genetic sex (Data-Field 22001), with sex chromosome aneuploidy (Data-Field 22019), heterozygosity outliers, or with variant missingness over 10%. To maximize statistical power, we excluded only participants who showed genetic relatedness of ten or more third-degree relatives (Data-Field 22021) and included participants of all ancestries (Data-Field 22006). Genotype arrays (Data-Field 22418) were lifted over from GRCh37 to GRCh38 using Picard LiftOverVCF. Imputation of lifted genotype array was performed by Genomics England (Data-Field 21008). Variant quality control was performed using plink2 (v2.0.0-a.6.9) and excluded variants with missingness greater than 10% across individuals, Hardy–Weinberg equilibrium *P*<1×10⁻⁶, minor allele frequency<1%, minor allele count<20, or multiallelic variants.

Functional annotation and gene mapping: We functionally annotated and mapped genes from GWAS results using the SNP2GENE pipeline as follows (FUMA v1.5.2) (Watanabe et al., 2017). Independent genome-wide significant SNPs were identified using a threshold of *P*<5×10⁻⁸ and pruned for linkage disequilibrium at *r²*<0.1 (1000 Genomes Phase 3, European panel). Genomic risk loci were defined by merging adjacent LD blocks within 250 kb. Additional candidate SNPs in LD with lead SNPs (*r²*≥0.6) were included for annotation. The extended MHC region was excluded from analysis. SNP-to-gene mapping was performed via three strategies: positional mapping of 35 kb upstream and 10 kb downstream, as commonly used in GWAS annotation and in Mullins et al. (2021), to allow inclusion of proximal regulatory variants near transcription start sites, expression quantitative trait loci (eQTL) mapping using brain-specific datasets (GTEx v8, PsychENCODE), and chromatin interaction mapping incorporating Hi-C and enhancer–promoter datasets from fetal and adult brain tissue. Only protein-coding genes were considered, based on Ensembl v102. Gene-based association was conducted using MAGMA v1.08, and gene expression enrichment was tested across GTEx v8 and BrainSpan transcriptomic datasets.

Pathway and tissue enrichment analysis: Genes mapped through positional, eQTL, or chromatin interaction methods were analysed using FUMA’s GENE2FUNC pipeline. Enrichment was tested using gene sets and GO terms from MSigDB and WikiPathways. Tissue specificity was evaluated using GTEx v8 expression data across 54 specific and 30 general tissue types, along with developmental brain expression profiles from BrainSpan.

Table S1. GWASs used for genetic correlation analysis

| Trait | Study | N |
| --- | --- | --- |
| ADHD | Demontis, Ditte, et al. "Genome-wide analyses of ADHD identify 27 risk loci, refine the genetic architecture and implicate several cognitive domains." Nature genetics 55.2 (2023): 198-208. | 225534 |
| Alzheimer | Bellenguez, Céline, et al. "New insights into the genetic etiology of Alzheimer’s disease and related dementias." Nature genetics 54.4 (2022): 412-436. | 1126563 |
| Anxiety | Otowa, Takeshi, et al. "Meta-analysis of genome-wide association studies of anxiety disorders." Molecular psychiatry 21.10 (2016): 1391-1399. | 18186 |
| Autism | Grove, Jakob, et al. "Identification of common genetic risk variants for autism spectrum disorder." *Nature genetics* 51.3 (2019): 431-444. | 46351 |
| BD | O’Connell, Kevin S., et al. "Genomics yields biological and phenotypic insights into bipolar disorder." *Nature* 639.8056 (2025): 968-975. | 840309 |
| MDD | Adams, Mark J., et al. "Trans-ancestry genome-wide study of depression identifies 697 associations implicating cell types and pharmacotherapies." *Cell* 188.3 (2025): 640-652. | 2000702 |
| OCD | Strom, Nora I., et al. "Genome-wide analyses identify 30 loci associated with obsessive–compulsive disorder." *Nature genetics* (2025): 1-13. | 1138106 |
| PTSD | Nievergelt, Caroline M., et al. "Genome-wide association analyses identify 95 risk loci and provide insights into the neurobiology of post-traumatic stress disorder." *Nature genetics* 56.5 (2024): 792-808. | 1222882 |
| SCZ | Trubetskoy, Vassily, et al. "Mapping genomic loci implicates genes and synaptic biology in schizophrenia." *Nature* 604.7906 (2022): 502-508. | 130644 |
| Tourette | Yu, Dongmei, et al. "Interrogating the genetic determinants of Tourette’s syndrome and other tic disorders through genome-wide association studies." *American Journal of Psychiatry* 176.3 (2019): 217-227. | 14307 |
| Neuroticism | Nagel, Mats, et al. "Meta-analysis of genome-wide association studies for neuroticism in 449,484 individuals identifies novel genetic loci and pathways." *Nature genetics* 50.7 (2018): 920-927. | 449484 |
| Guilt | http://www.nealelab.is/uk-biobank/ | 351907 |
| Depressive episode onset | http://www.nealelab.is/uk-biobank/ | 61033 |
| Cognitive processing accuracy | Li, Mingyang, et al. "Cognitive processing speed and accuracy are intrinsically different in genetic architecture and brain phenotypes." *Nature Communications* 15.1 (2024): 7786. | 1783727 |
| Reaction time | Davies, Gail, et al. "Study of 300,486 individuals identifies 148 independent genetic loci influencing general cognitive function." *Nature communications* 9.1 (2018): 2098. | 330069 |
| Reasoning | Davies, Gail, et al. "Genome-wide association study of cognitive functions and educational attainment in UK Biobank (N= 112 151)." *Molecular psychiatry* 21.6 (2016): 758-767. | 36035 |
| SC and FC of RSNs | Tissink, Elleke, et al. "The genetic architectures of functional and structural connectivity properties within cerebral resting-state networks." *eneuro* 10.4 (2023). | 24,336 |

Traits included in the genetic correlation analysis, with corresponding study citation and sample size (N). ADHD = attention-deficit/hyperactivity disorder, AD = Alzheimer disease, ASD = autism spectrum disorder, BD = bipolar disorder, MDD = major depressive disorder, OCD = obsessive–compulsive disorder, PTSD = post-traumatic stress disorder, SCZ = schizophrenia, TS = Tourette syndrome.

### RESULTS

#### Regional Structure-Function Coupling:

Figure S1. Between-network regional SC-FC coupling across cortical and subcortical regions

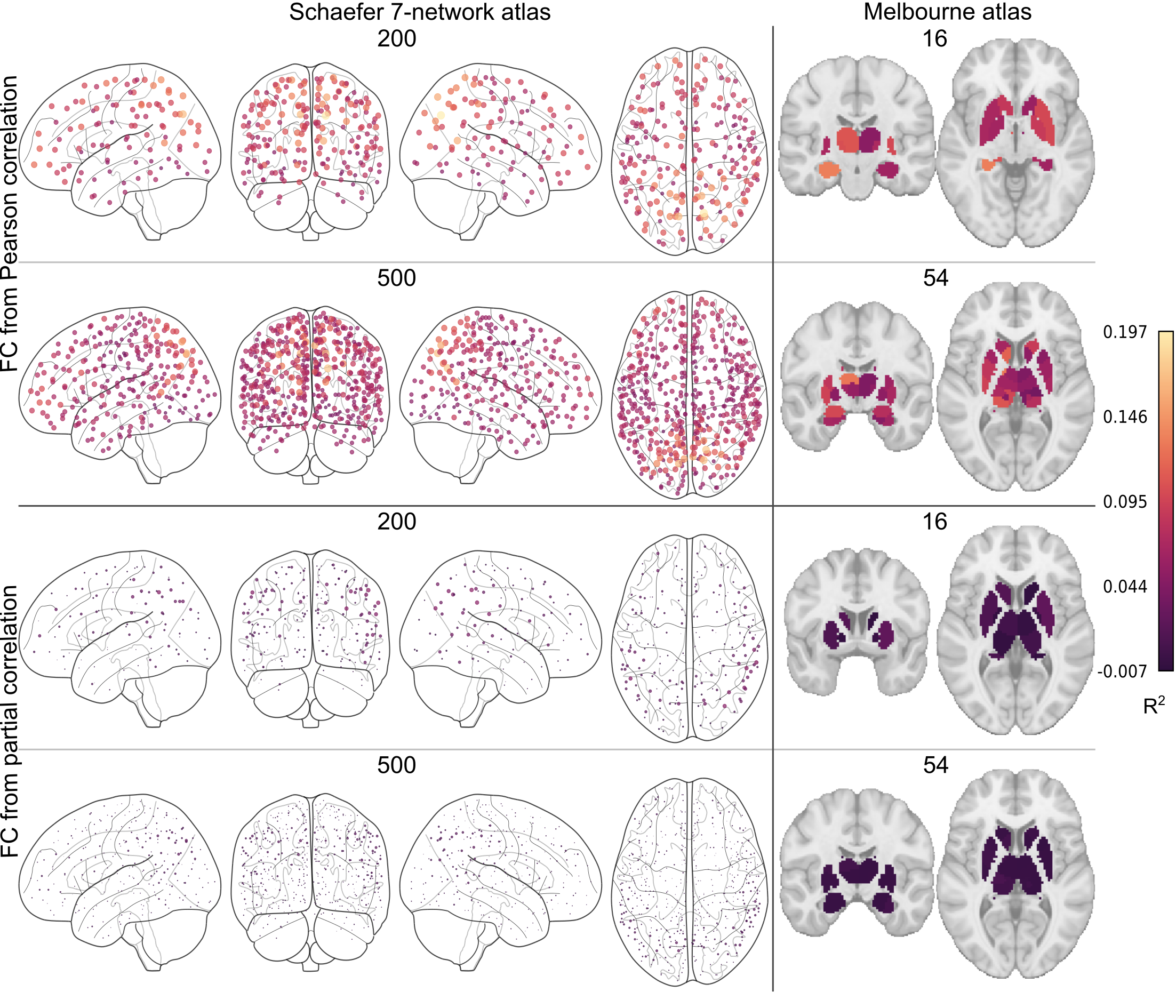

Between-network structure-function coupling (R²) was computed for 200 or 500 cortical and 16 or 54 subcortical regions, with FC estimated using Pearson or partial correlations. Larger and lighter dots indicate stronger coupling. FC = functional connectivity.

Figure S2. Within-network regional SC-FC coupling across cortical and subcortical regions

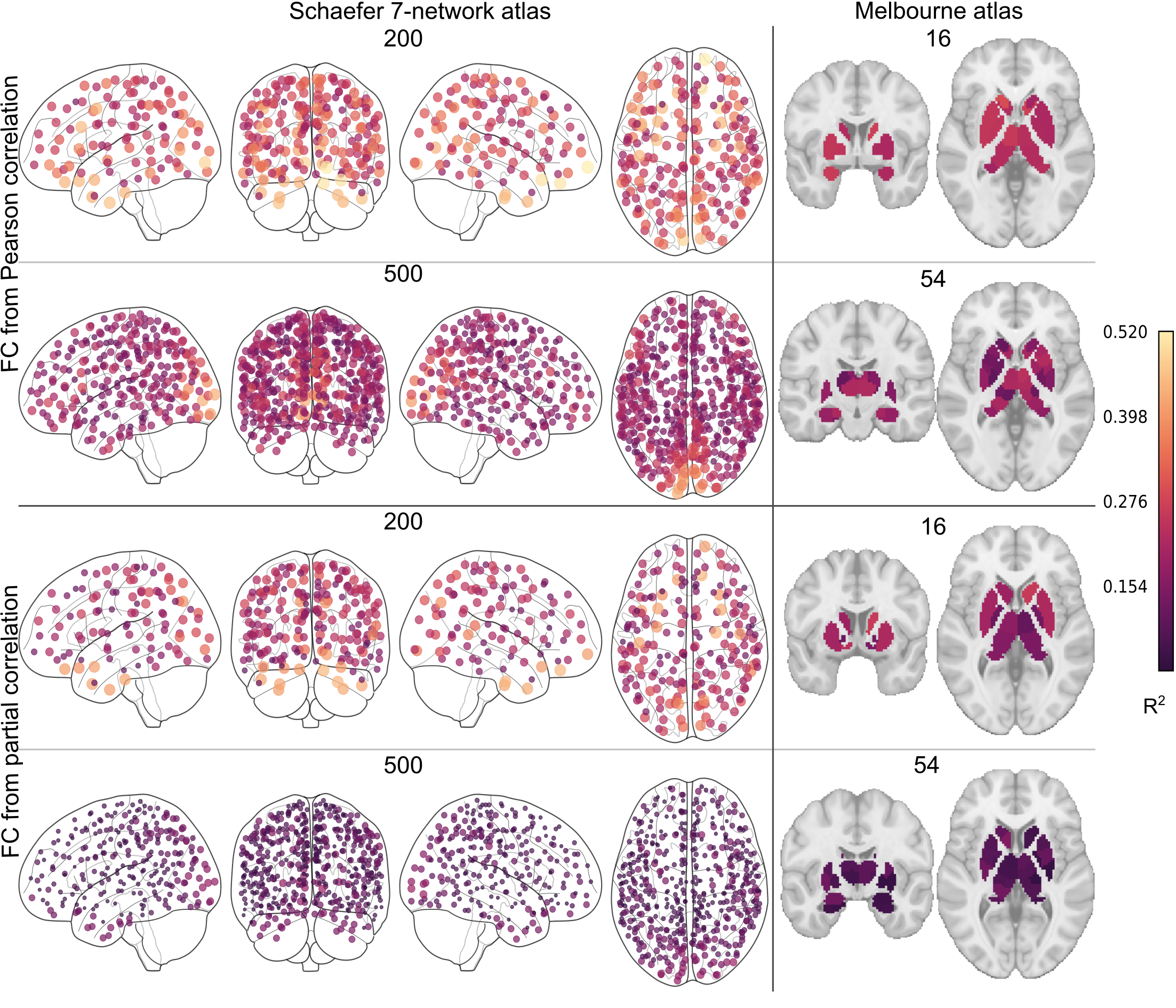

Within-network structure-function coupling (R²) was computed for 200 or 500 cortical and 16 or 54 subcortical regions, with FC estimated using Pearson or partial correlations. Larger and lighter dots indicate stronger coupling. FC = functional connectivity.

Table S2. Regions with the highest between-network SC-FC coupling

| Region | R² Mean (SD) | R² Range | Centroid coordinates |
| --- | --- | --- | --- |
| 216 regions, Pearson | | | |
| Right precuneus cortex | 0.197 (0.11) | -0.00–0.51 | 14, -70, 37 |
| Left precuneus cortex | 0.168 (0.10) | -0.01–0.47 | -9, -73, 38 |
| Right precuneus cortex | 0.161 (0.10) | -0.01–0.46 | 12, -55, 15 |
| Right superior lateral occipital cortex | 0.158 (0.08) | -0.00–0.45 | 15, -73, 53 |
| Right superior parietal lobule | 0.144 (0.07) | -0.01–0.39 | 21, -48, 70 |
| Left superior parietal lobule | 0.141 (0.06) | -0.00–0.40 | -17, -53, 68 |
| Right superior lateral occipital cortex | 0.139 (0.07) | -0.01–0.42 | 33, -75, 32 |
| Right precuneous cortex | 0.138 (0.07) | -0.01–0.34 | 11, -36, 47 |
| 216 regions, partial | | | |
| Right posterior supramarginal gyrus | 0.048 (0.05) | -0.02–0.27 | 53, -42, 48 |
| Right posterior supramarginal gyrus | 0.047 (0.03) | -0.02–0.16 | 62, -37, 37 |
| Right superior lateral occipital cortex | 0.045 (0.04) | -0.02–0.19 | 33, -75, 32 |
| Left posterior supramarginal gyrus | 0.042 (0.04) | -0.02–0.22 | -53, -51, 46 |
| Right superior lateral occipital cortex | 0.041 (0.04) | -0.02–0.22 | 37, -63, 47 |
| Right anterior supramarginal gyrus | 0.040 (0.03) | -0.02–0.16 | 60, -26, 27 |
| Right superior lateral occipital cortex | 0.039 (0.04) | -0.02–0.29 | 51, -59, 44 |
| Right middle temporooccipital gyrus | 0.035 (0.03) | -0.02–0.16 | 57, -45, 9 |
| 554 regions, Pearson | | | |
| Right precuneus cortex | 0.175 (0.09) | -0.01–0.43 | 16, -64, 29 |
| Right superior lateral occipital cortex | 0.170 (0.09) | -0.00–0.41 | 14, -73, 42 |
| Left precuneus cortex | 0.165 (0.08) | 0.01–0.41 | -9, -76, 42 |
| Right precuneus cortex | 0.158 (0.08) | 0.02–0.48 | 4, -70, 49 |
| Left precuneus cortex | 0.151 (0.08) | 0.00–0.38 | -4, -72, 52 |
| Left superior lateral occipital cortex | 0.140 (0.07) | 0.00–0.33 | -29, -76, 41 |
| Right precuneus cortex | 0.136 (0.08) | 0.01–0.40 | 14, -56, 17 |
| Right superior lateral occipital cortex | 0.133 (0.07) | -0.00–0.42 | 30, -68, 33 |
| 554 regions, partial | | | |
| Left precuneus | 0.021 (0.02) | -0.01–0.11 | -9, -76, 42 |
| Right superior lateral occipital cortex | 0.019 (0.02) | -0.01–0.08 | 47, -71, 25 |
| Left precuneus cortex | 0.018 (0.02) | -0.01–0.12 | -11, -69, 30 |
| Right angular gyrus | 0.017 (0.02) | -0.01–0.09 | 41, -56, 48 |
| Right posterior supramarginal gyrus | 0.017 (0.02) | -0.01–0.10 | 60, -40, 36 |
| Left superior lateral occipital cortex | 0.016 (0.02) | -0.01–0.12 | -29, -76, 41 |
| Left posterior supramarginal gyrus | 0.016 (0.01) | -0.01–0.07 | -53, -50, 46 |
| Right anterior supramarginal gyrus | 0.016 (0.02) | -0.01–0.11 | 61, -33, 46 |

Values reflect mean (SD) and range of adjusted R² estimates of between-network structure-function coupling across all participants. “216 regions, Pearson”; “216 regions, partial”; “554 regions, Pearson”; and “554 regions, partial” refer to the parcellation resolution (Schaefer cortical + Melbourne subcortical atlases) and the functional connectivity method (Pearson or partial correlation). Centroid coordinates are reported in MNI152 space. SD = standard deviation.

Table S3. Regions with the lowest between-network SC-FC coupling

| Region | R² Mean (SD) | R² Range | Centroid coordinates |
| --- | --- | --- | --- |
| 216 regions, Pearson | | | |
| Right posterior parahippocampal gyrus | 0.033 (0.04) | -0.02-0.23 | 28, -36, -14 |
| Left lingual gyrus | 0.037 (0.04) | -0.02-0.25 | -24, -53, -9 |
| Right anterior cingulate gyrus | 0.038 (0.05) | -0.02-0.31 | 7, 9, 41 |
| Right anterior thalamus | 0.041 (0.04) | -0.02-0.25 | 10, -14, 8 |
| Right insular cortex | 0.042 (0.05) | -0.02-0.28 | 41, 6, -15 |
| Right hippocampus | 0.042 (0.04) | -0.02-0.25 | 28, -22, -14 |
| Right occipital fusiform gyrus | 0.043 (0.05) | -0.02-0.27 | 29, -69, -12 |
| Left insular cortex | 0.044 (0.05) | -0.02-0.41 | -33, 20, 5 |
| 216 regions, partial | | | |
| Left Lingual gyrus | -0.007 (0.01) | -0.02–0.06 | -10, -67, -4 |
| Left Accumbens | -0.007 (0.01) | -0.02–0.04 | -10, 14, -6 |
| Right Occipital pole | -0.007 (0.01) | -0.02–0.04 | 11, -92, -5 |
| Right Accumbens | -0.007 (0.01) | -0.02–0.04 | 12, 14, -6 |
| Left Paracingulate gyrus | -0.006 (0.01) | -0.02–0.05 | -6, 36, -10 |
| Right Lingual gyrus | -0.006 (0.01) | -0.02–0.05 | 12, -65, -5 |
| Left Pallidum | -0.006 (0.01) | -0.02–0.05 | -18, -4, -2 |
| Left Occipital pole | -0.006 (0.01) | -0.02–0.04 | -27, -95, -12 |
| 554 regions, Pearson | | | |
| Right lingual gyrus | 0.029 (0.03) | -0.01–0.25 | 17, -43, -3 |
| Right anterior cingulate gyrus | 0.029 (0.03) | -0.01–0.20 | 7, 8, 41 |
| Left temporal occipital fusiform cortex | 0.030 (0.03) | -0.01–0.15 | -25, -57, -10 |
| Left lingual gyrus | 0.030 (0.03) | -0.01–0.16 | -23, -48, -6 |
| Right lingual gyrus | 0.032 (0.03) | -0.01–0.16 | 14, -58, -5 |
| Left posterior parahippocampal gyrus | 0.032 (0.03) | -0.01–0.20 | -29, -37, -15 |
| Right visual area 5 | 0.032 (0.03) | -0.01–0.15 | 27, -53, -9 |
| Left temporal occipital fusiform cortex | 0.033 (0.03) | -0.00–0.17 | -35, -60, -17 |
| 554 regions, partial | | | |
| Right ventroanterior caudate | -0.003 (0.00) | -0.01–0.02 | 10, 12, 4 |
| Left nucleus accumbens shell | -0.003 (0.00) | -0.01–0.02 | -10, 10, -6 |
| Right nucleus accumbens core | -0.003 (0.00) | -0.01–0.02 | 14, 18, -2 |
| Right postcentral gyrus | -0.002 (0.01) | -0.01–0.02 | 53, -10, 36 |
| Right frontal orbital cortex | -0.002 (0.00) | -0.01–0.02 | 12, 25, -21 |
| Right posterior temporal fusiform cortex | -0.002 (0.00) | -0.01–0.02 | 40, -15, -31 |
| Right posterior inferior temporal gyrus | -0.002 (0.00) | -0.01–0.02 | 53, -17, -33 |
| Right posterior middle temporal gyrus | -0.002 (0.00) | -0.01–0.02 | 58, -8, -27 |

Values reflect mean (SD) and range of adjusted R² estimates of between-network structure-function coupling across all participants. “216 regions, Pearson”; “216 regions, partial”; “554 regions, Pearson”; and “554 regions, partial” refer to the parcellation resolution (Schaefer cortical + Melbourne subcortical atlases) and the functional connectivity method (Pearson or partial correlation). Centroid coordinates are reported in MNI152 space. SD = standard deviation.

Table S4. Regions with the highest within-network structure-function coupling

| Region | R² Mean (SD) | R² Range | Centroid coordinates |
| --- | --- | --- | --- |
| 216 regions, Pearson | | | |
| Right frontal pole | 0.520 (0.21) | 0.02–0.95 | 15, 64, -8 |
| Right frontal pole | 0.506 (0.21) | 0.05–0.97 | 12, 39, -22 |
| Right frontal orbital cortex | 0.503 (0.21) | 0.00–0.95 | 28, 22, -19 |
| Left occipital pole | 0.486 (0.17) | 0.03–0.88 | -5, -93, -4 |
| Right posterior inferior temporal gyrus | 0.462 (0.20) | 0.04–0.93 | 47, -12, -35 |
| Left anterior temporal fusiform cortex | 0.459 (0.20) | 0.04–0.93 | -29, -6, -39 |
| Left middle frontal gyrus | 0.451 (0.13) | 0.08–0.80 | -44, 20, 27 |
| 216 regions, partial | | | |
| Right frontal pole | 0.446 (0.21) | 0.02–0.97 | 12, 39, -22 |
| Right anterior parahippocampal gyrus | 0.444 (0.20) | 0.03–0.90 | 25, -11, -32 |
| Right temporal pole | 0.427 (0.19) | 0.02–0.89 | 30, 9, -38 |
| Right posterior inferior temporal gyrus | 0.425 (0.20) | 0.02–0.96 | 47, -12, -35 |
| Left posterior inferior temporal gyrus | 0.419 (0.19) | 0.03–0.89 | -45, -20, -30 |
| Left temporal pole | 0.417 (0.21) | 0.02–0.91 | -28, 10, -34 |
| Left temporal pole | 0.412 (0.19) | 0.02–0.91 | -43, 8, -19 |
| 554 regions, Pearson | | | |
| Left occipital pole | 0.434 (0.14) | 0.02–0.79 | -10, -102, 6 |
| Left occipital pole | 0.431 (0.14) | 0.07–0.81 | -5, -91, 1 |
| Left occipital pole | 0.419 (0.15) | 0.03–0.76 | -9, -98, -11 |
| Right occipital pole | 0.407 (0.14) | 0.04–0.74 | 11, -97, 0 |
| Right intracalcarine cortex | 0.395 (0.14) | 0.06–0.73 | 7, -84, 5 |
| Left intracalcarine cortex | 0.381 (0.14) | 0.03–0.75 | -6, -78, 11 |
| Left occipital pole | 0.379 (0.13) | 0.04–0.73 | -6, -95, 17 |
| Left cuneal cortex | 0.363 (0.13) | 0.04–0.70 | -3, -84, 25 |
| 554 regions, Partial | | | |
| Right occipital pole | 0.190 (0.08) | 0.01–0.45 | 11, -97, 0 |
| Right inferior lateral occipital cortex | 0.185 (0.10) | 0.01–0.46 | 41, -86, -5 |
| Left occipital pole | 0.181 (0.08) | 0.02–0.51 | -5, -91, 1 |
| Left occipital pole | 0.180 (0.09) | 0.00–0.44 | -25, -96, 5 |
| Left occipital pole | 0.174 (0.09) | 0.02–0.47 | -10, -102, 6 |
| Right occipital pole | 0.173 (0.09) | 0.00–0.51 | 26, -97, 6 |
| Left occipital pole | 0.169 (0.08) | 0.00–0.44 | -9, -98, -11 |
| Left intracalcarine cortex | 0.169 (0.08) | 0.01–0.43 | -6, -78, 11 |

Values reflect mean (SD) and range of adjusted R² estimates of within-network structure-function coupling across all participants. “216 regions, Pearson”; “216 regions, partial”; “554 regions, Pearson”; and “554 regions, partial” refer to the parcellation resolution (Schaefer cortical + Melbourne subcortical atlases) and the functional connectivity method (Pearson or partial correlation). Centroid coordinates are reported in MNI152 space. SD = standard deviation.

Table S5. Regions with the lowest within-network structure-function coupling

| Region | R² Mean (SD) | R² Range | Centroid coordinates |
| --- | --- | --- | --- |
| 216 regions, Pearson | | | |
| Left superior frontal gyrus | 0.127 (0.08) | 0.00–0.50 | -24, 25, 49 |
| Right posterior parahippocampal gyrus | 0.149 (0.11) | 0.01–0.66 | 28, -36, -14 |
| Right temporal pole | 0.154 (0.09) | 0.01–0.56 | 47, 13, -30 |
| Right middle frontal gyrus | 0.155 (0.09) | 0.00–0.49 | 29, 30, 42 |
| Right superior frontal gyrus | 0.166 (0.09) | 0.01–0.50 | 23, 24, 53 |
| Right occipital fusiform gyrus | 0.170 (0.10) | 0.01–0.64 | 29, -69, -12 |
| Left superior lateral occipital cortex | 0.172 (0.09) | 0.01–0.47 | -46, -66, 38 |
| Left posterior parahippocampal gyrus | 0.173 (0.09) | 0.01–0.45 | -26, -32, -18 |
| 216 regions, partial | | | |
| Right postcentral gyrus | 0.089 (0.06) | 0.00–0.34 | 58, -5, 31 |
| Left posterior parahippocampal gyrus | 0.093 (0.06) | 0.00–0.33 | -26, -32, -18 |
| Right anterior cingulate gyrus | 0.098 (0.07) | 0.00–0.39 | 6, 29, 15 |
| Left posterior middle temporal gyrus | 0.099 (0.06) | 0.00–0.46 | -60, -19, -22 |
| Left superior frontal gyrus | 0.102 (0.06) | 0.00–0.33 | -24, 25, 49 |
| Right inferior frontal cortex pars triangularis | 0.106 (0.07) | 0.01–0.33 | 51, 28, 0 |
| Left frontal orbital cortex | 0.108 (0.07) | 0.00–0.36 | -35, 20, -13 |
| Left posterior cingulate gyrus | 0.110 (0.07) | 0.01–0.39 | -4, -31, 36 |
| 554 regions, Pearson | | | |
| Right anterior middle temporal gyrus | 0.089 (0.06) | 0.00–0.28 | 60, -5, -16 |
| Right PFCdPFCm 11 | 0.091 (0.06) | 0.00–0.35 | 22, 40, 37 |
| Left posterior parahippocampal gyrus | 0.096 (0.07) | 0.00–0.35 | -19, -35, -14 |
| Left posterior globus pallidus | 0.096 (0.06) | 0.00–0.36 | -22, -8, -2 |
| Right posterior globus pallidus | 0.097 (0.06) | 0.01–0.34 | 24, -8, -2 |
| Left anterior globus pallidus | 0.100 (0.06) | 0.00–0.44 | -16, 0, -2 |
| Right ventroanterior putamen | 0.102 (0.06) | 0.00–0.34 | 22, 12, -6 |
| Right PFCdPFCm 13 | 0.102 (0.06) | 0.00–0.32 | 25, 26, 46 |
| 554 regions, partial | | | |
| Left frontal orbital cortex | 0.032 (0.02) | 0.00–0.14 | -29, 17, -17 |
| Left anterior cingulate gyrus | 0.033 (0.02) | 0.00–0.17 | -3, 26, 19 |
| Left temporal region 3 | 0.034 (0.02) | 0.00–0.13 | -60, -36, -18 |
| Right PFCdPFCm 2 | 0.038 (0.03) | 0.00–0.19 | 6, 35, -6 |
| Right frontal pole | 0.038 (0.03) | 0.00–0.15 | 35, 37, -13 |
| Left posterior parahippocampal gyrus | 0.038 (0.03) | 0.00–0.18 | -20, -25, -22 |
| Left posterior parahippocampal gyrus | 0.038 (0.03) | 0.00–0.16 | -19, -35, -14 |
| Left posterior cingulate cortex | 0.038 (0.02) | 0.00–0.15 | -2, -16, 37 |

Values reflect mean (SD) and range of adjusted R² estimates of within-network structure-function coupling across all participants. “216 regions, Pearson”; “216 regions, partial”; “554 regions, Pearson”; and “554 regions, partial” refer to the parcellation resolution (Schaefer cortical + Melbourne subcortical atlases) and the functional connectivity method (Pearson or partial correlation). Centroid coordinates are reported in MNI152 space. SD = standard deviation.

#### Structure-Function Coupling Associations with Degree and Strength:

Table S6. Associations of structure-function coupling with degree and strength

|  |  | 216 regions | | 554 regions | |
| --- | --- | --- | --- | --- | --- |
|  | FC type | R² | p value | R² | p value |
| Structural degree | Pearson | 0.05 | 0.001 | 0.03 | <0.001 |
|  | Partial | 0.01 | 0.084 | 0.00 | 0.175 |
| Structural strength | Pearson | 0.01 | 0.220 | 0.04 | <0.001 |
|  | Partial | 0.01 | 0.135 | 0.08 | <0.001 |
| Functional strength | Pearson | 0.15 | <0.001 | 0.25 | <0.001 |
|  | Partial | 0.32 | <0.001 | 0.00 | 0.709 |

Values show the variance in regional structure-function coupling explained by structural degree, structural strength, and functional strength for each functional connectivity type at the 216 and 554 resolutions.

Figure S3. Associations between regional structure-function coupling and degree and strength when functional connectivity was calculated with Pearson correlation

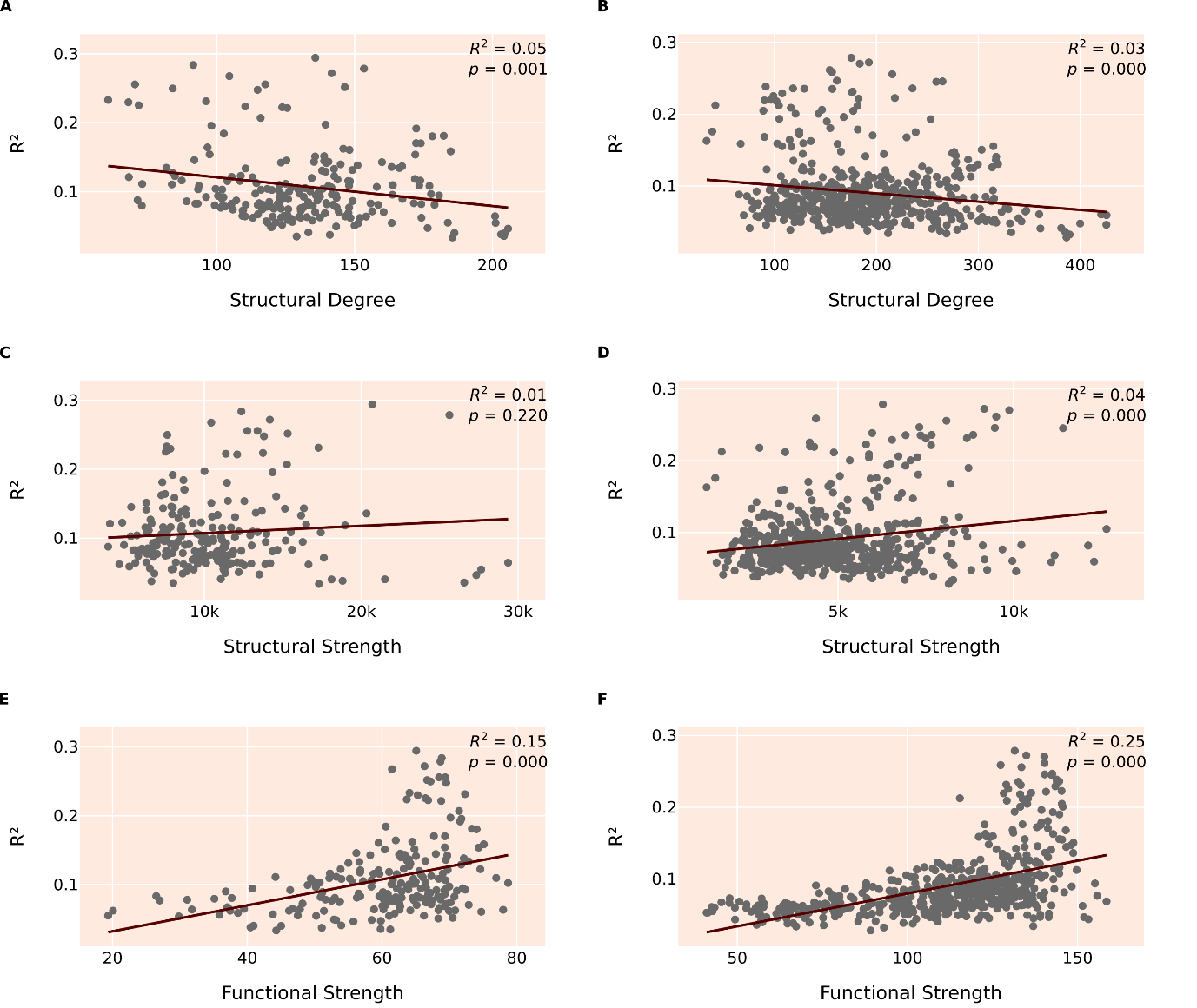

Linear regression of SFC in the 216-region parcellation with (A) structural degree, (C) structural strength, (E) functional strength, and in the 554-region parcellation with (B) structural degree, (D) structural strength, (F) functional strength.

Figure S4. Associations between regional structure-function coupling and degree and strength when functional connectivity was calculated with partial correlation

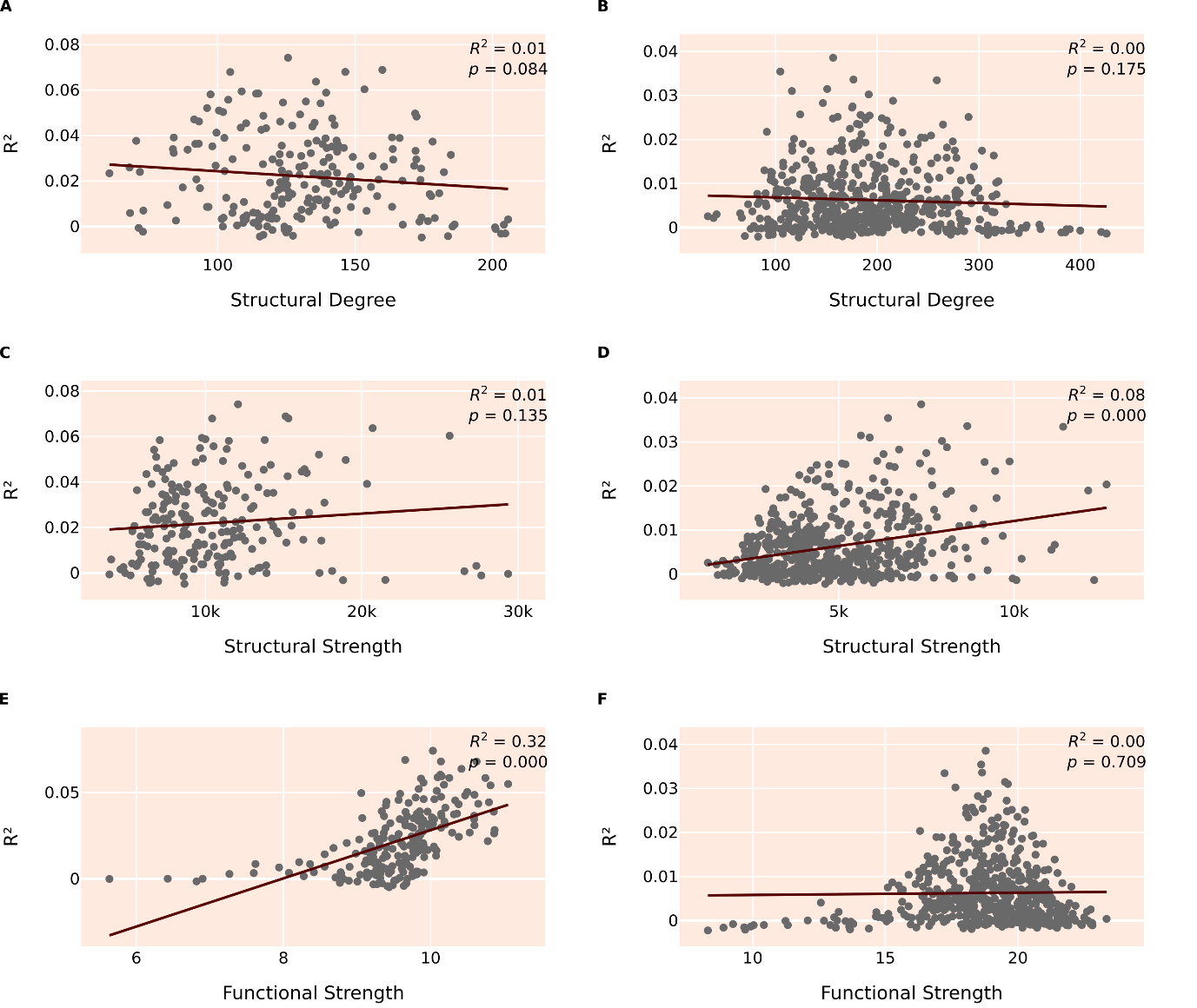

Linear regression of SFC in the 216-region parcellation with (A) structural degree, (C) structural strength, (E) functional strength, and in the 554-region parcellation with (B) structural degree, (D) structural strength, (F) functional strength.

#### Structure-Function Coupling of Resting-State Networks:

Table S7. Resting-state networks SC-FC coupling

| Network | R^2^ Mean (SD) | R^2^ Range |
| --- | --- | --- |
|  | 216 regions, Pearson |  |
| Vis | 0.207 (0.06) | 0.08-0.29 |
| SMN | 0.107 (0.03) | 0.06-0.16 |
| DAN | 0.113 (0.04) | 0.05-0.19 |
| VAN | 0.077 (0.03) | 0.04-0.14 |
| Lim | 0.073 (0.01) | 0.06-0.10 |
| CEN | 0.089 (0.02) | 0.06-0.16 |
| DMN | 0.095 (0.02) | 0.04-0.15 |
| Sub | 0.059 (0.02) | 0.03-0.11 |
| 216 regions, partial | | |
| Vis | 0.039 (0.021) | 0.000-0.074 |
| SMN | 0.022 (0.017) | -0.001-0.059 |
| DAN | 0.026 (0.011) | 0.007-0.048 |
| VAN | 0.017 (0.015) | 0.001-0.051 |
| Lim | 0.002 (0.004) | -0.002-0.011 |
| CEN | 0.024 (0.017) | -0.004-0.058 |
| DMN | 0.023 (0.015) | -0.002-0.069 |
| Sub | -0.001 (0.003) | -0.005-.007 |
| 554 regions, Pearson | | |
| Vis | 0.179 (0.056) | 0.049-0.279 |
| SMN | 0.087 (0.018) | 0.045-0.131 |
| DAN | 0.092 (0.032) | 0.044-0.156 |
| VAN | 0.064 (0.016) | 0.034-0.108 |
| Lim | 0.062 (0.008) | 0.051-0.086 |
| CEN | 0.082 (0.025) | 0.038-0.160 |
| DMN | 0.080 (0.018) | 0.044-0.141 |
| Sub | 0.054 (0.013) | 0.028-0.091 |
| 554 regions, partial | | |
| Vis | 0.013 (0.010) | -0.001–0.039 |
| SMN | 0.006 (0.006) | -0.000–0.031 |
| DAN | 0.007 (0.004) | 0.001–0.018 |
| VAN | 0.004 (0.005) | -0.001–0.022 |
| Lim | -0.001 (0.001) | -0.002–0.004 |
| CEN | 0.008 (0.007) | -0.001–0.025 |
| DMN | 0.007 (0.006) | -0.001–0.025 |
| Sub | -0.001 (0.001) | -0.002–0.002 |

Values reflect mean (SD) and range of adjusted R² estimates of structure-function coupling within each resting-state network. “216” and “554” refer to the parcellation resolutions derived from the Schaefer cortical atlas (200- or 500-region versions) combined with the Melbourne subcortical atlas (16 or 54 regions, respectively). Pearson and partial denote the functional connectivity estimation method. CEN = central executive; DAN = dorsal attention; DMN = default mode; Lim = limbic; SMN = sensorimotor; Sub = subcortical; VAN = ventral attention; Vis = visual.

Figure S5. Between-network regional structure-function coupling (R^2^) averaged across resting-state networks

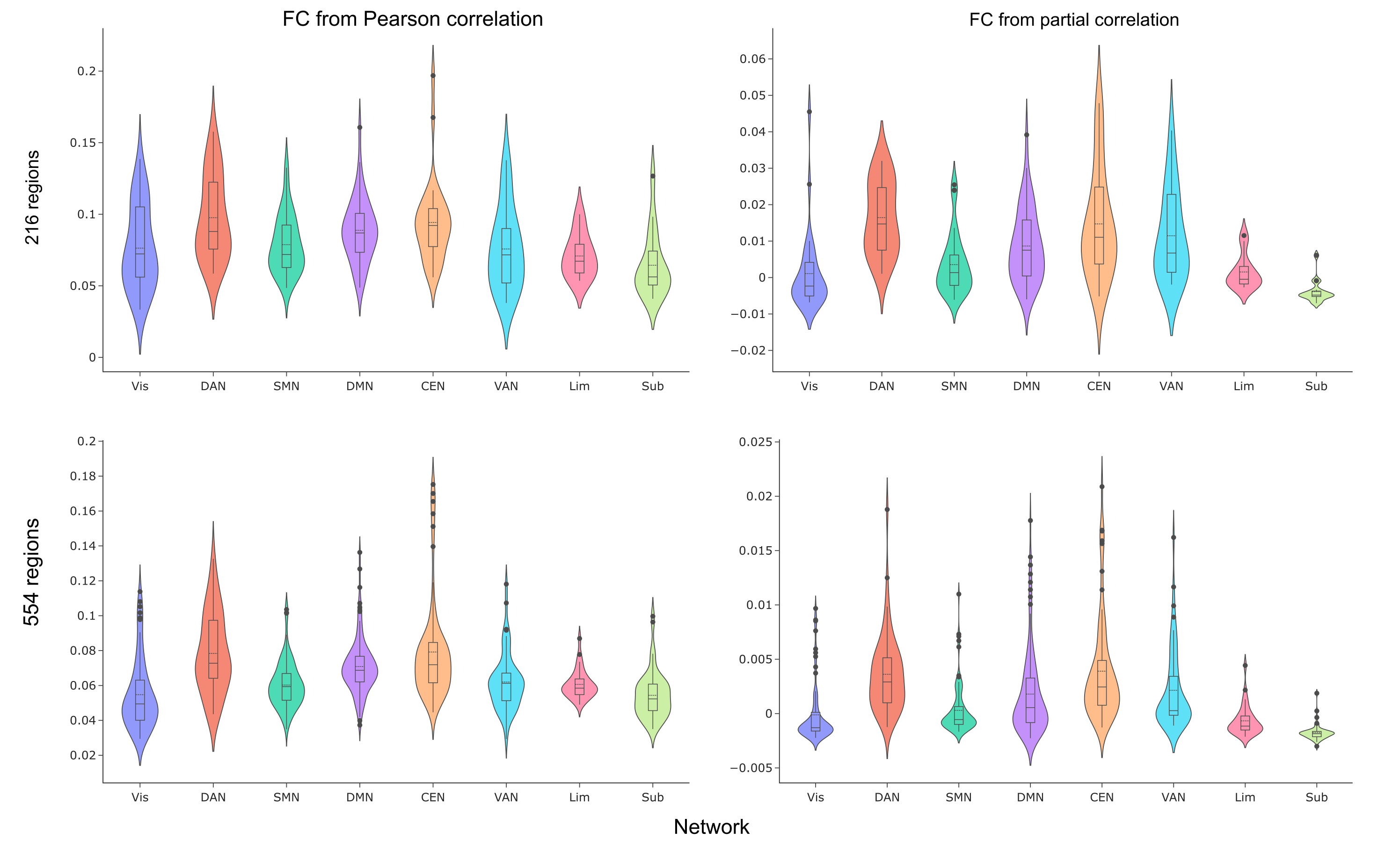

Between-network structure-function coupling (R^2^) was averaged across resting-state networks as assigned by the 200 or 500-region Schaefer 7-network atlas, and with 16 or 54 subcortical regions in the Melbourne atlas, with functional connectivity calculated using Pearson or partial correlation. CEN = central executive; DAN = dorsal attention; DMN = default mode; FC = functional connectivity; Lim = limbic; SMN = sensorimotor; Sub = subcortical; VAN = ventral attention; Vis = visual.

Figure S6. Within-network regional structure-function coupling (R^2^) averaged across resting-state networks

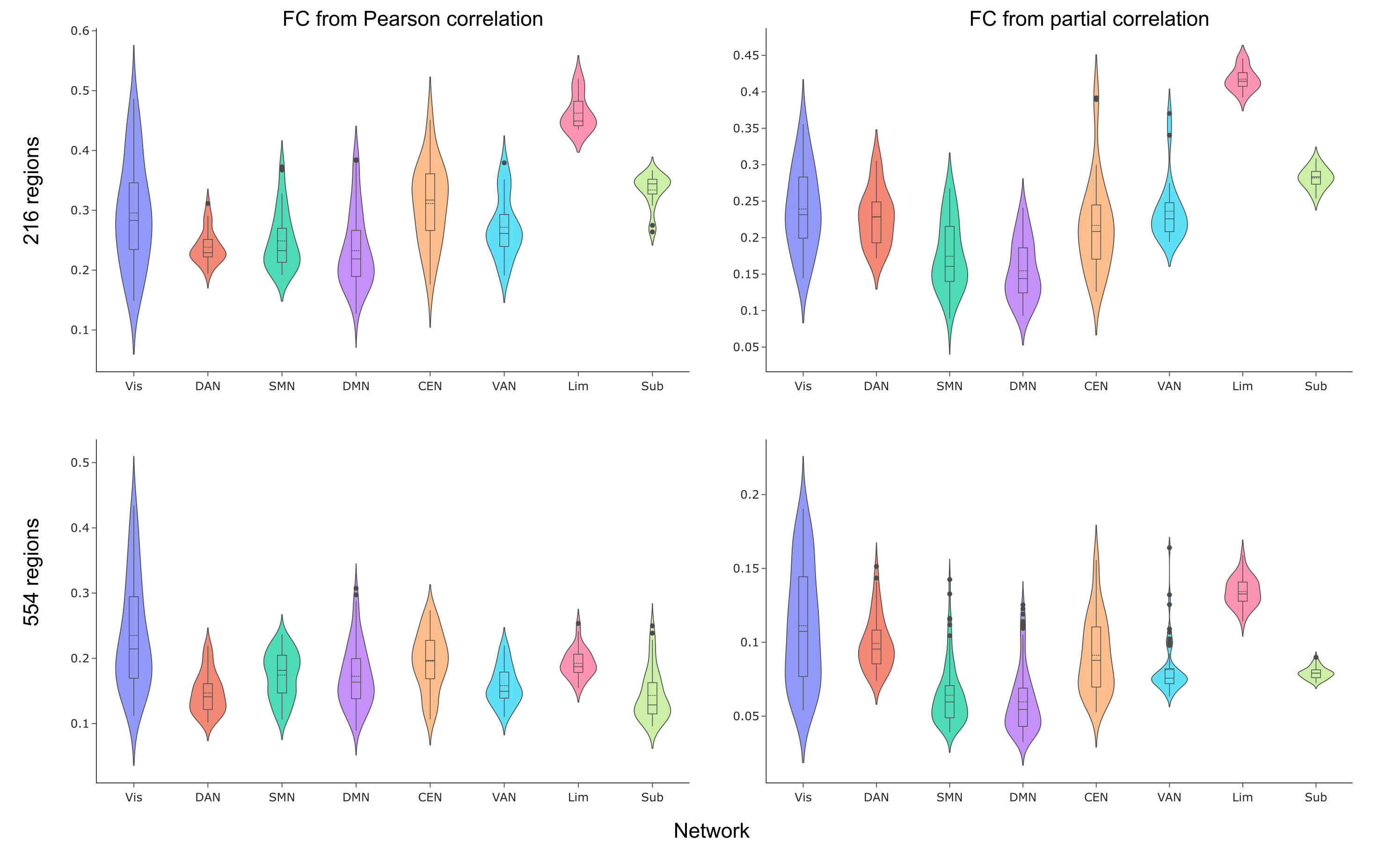

Within-network structure-function coupling (R^2^) was averaged across resting-state networks as assigned by the 200 or 500-region Schaefer 7-network atlas, and with 16 or 54 subcortical regions in the Melbourne atlas, with functional connectivity calculated using Pearson or partial correlation. CEN = central executive; DAN = dorsal attention; DMN = default mode; FC = functional connectivity; Lim = limbic; SMN = sensorimotor; Sub = subcortical; VAN = ventral attention; Vis = visual.

Table S8. Resting-state networks between-network SC-FC coupling

| Network | R^2^ Mean (SD) | R^2^ Range |
| --- | --- | --- |
|  | 216 regions, Pearson |  |
| Vis | 0.207 (0.06) | 0.08-0.29 |
| SMN | 0.107 (0.03) | 0.06-0.16 |
| DAN | 0.113 (0.04) | 0.05-0.19 |
| VAN | 0.077 (0.03) | 0.04-0.14 |
| Lim | 0.073 (0.01) | 0.06-0.10 |
| CEN | 0.089 (0.02) | 0.06-0.16 |
| DMN | 0.095 (0.02) | 0.04-0.15 |
| Sub | 0.059 (0.02) | 0.03-0.11 |
| 216 regions, partial | | |
| Vis | 0.039 (0.021) | 0.000-0.074 |
| SMN | 0.022 (0.017) | -0.001-0.059 |
| DAN | 0.026 (0.011) | 0.007-0.048 |
| VAN | 0.017 (0.015) | 0.001-0.051 |
| Lim | 0.002 (0.004) | -0.002-0.011 |
| CEN | 0.024 (0.017) | -0.004-0.058 |
| DMN | 0.023 (0.015) | -0.002-0.069 |
| Sub | -0.001 (0.003) | -0.005-.007 |
| 554 regions, Pearson | | |
| Vis | 0.055 (0.020) | 0.029–0.114 |
| SMN | 0.060 (0.013) | 0.035–0.104 |
| DAN | 0.078 (0.023) | 0.044–0.133 |
| VAN | 0.062 (0.017) | 0.029–0.118 |
| Lim | 0.061 (0.008) | 0.049–0.087 |
| CEN | 0.079 (0.029) | 0.045–0.175 |
| DMN | 0.071 (0.016) | 0.037–0.136 |
| Sub | 0.054 (0.013) | 0.035–0.100 |
| 554 regions, partial | | |
| Vis | -0.000 (0.003) | -0.002–0.010 |
| SMN | 0.000 (0.002) | -0.002–0.011 |
| DAN | 0.004 (0.004) | -0.001–0.019 |
| VAN | 0.002 (0.004) | -0.001–0.016 |
| Lim | -0.001 (0.001) | -0.002–0.004 |
| CEN | 0.004 (0.005) | -0.001–0.021 |
| DMN | 0.002 (0.004) | -0.002–0.018 |
| Sub | -0.002 (0.001) | -0.003–0.002 |

Values reflect mean (SD) and range of adjusted R² estimates of structure-function coupling within each resting-state network. “216” and “554” refer to the parcellation resolutions derived from the Schaefer cortical atlas (200- or 500-region versions) combined with the Melbourne subcortical atlas (16 or 54 regions, respectively). Pearson and partial denote the functional connectivity estimation method. CEN = central executive; DAN = dorsal attention; DMN = default mode; Lim = limbic; SMN = sensorimotor; Sub = subcortical; VAN = ventral attention; Vis = visual.

Table S9. Resting-state networks within-network structure-function coupling

| Network | R^2^ Mean (SD) | R^2^ Range |
| --- | --- | --- |
|  | 216 regions, Pearson |  |
| Vis | 0.295 (0.089) | 0.15-0.49 |
| SMN | 0.249 (0.049) | 0.19-0.37 |
| DAN | 0.238 (0.028) | 0.19-0.31 |
| VAN | 0.272 (0.051) | 0.19-0.38 |
| Lim | 0.462 (0.030) | 0.44-0.52 |
| CEN | 0.311 (0.067) | 0.18-0.45 |
| DMN | 0.233 (0.064) | 0.13-0.38 |
| Sub | 0.334 (0.029) | 0.26-0.37 |
| 216 regions, partial | | |
| Vis | 0.239 (0.057) | 0.15-0.36 |
| SMN | 0.175 (0.047) | 0.09-0.27 |
| DAN | 0.228 (0.039) | 0.17-0.31 |
| VAN | 0.236 (0.044) | 0.19-0.37 |
| Lim | 0.417 (0.016) | 0.39-0.45 |
| CEN | 0.217 (0.065) | 0.13-0.39 |
| DMN | 0.155 (0.041) | 0.09-0.24 |
| Sub | 0.282 (0.014) | 0.25-0.31 |
| 554 regions, Pearson | | |
| Vis | 0.235 (0.084) | 0.11-0.43 |
| SMN | 0.174 (0.036) | 0.11-0.24 |
| DAN | 0.147 (0.031) | 0.10-0.22 |
| VAN | 0.158 (0.029) | 0.11-0.22 |
| Lim | 0.192 (0.022) | 0.16-0.25 |
| CEN | 0.196 (0.045) | 0.11-0.27 |
| DMN | 0.173 (0.048) | 0.09-0.31 |
| Sub | 0.143 (0.038) | 0.10-0.25 |
| 554 regions, partial | | |
| Vis | 0.111 (0.040) | 0.05-0.19 |
| SMN | 0.064 (0.021) | 0.04-0.14 |
| DAN | 0.099 (0.018) | 0.07-0.15 |
| VAN | 0.081 (0.018) | 0.06-0.16 |
| Lim | 0.134 (0.010) | 0.11-0.16 |
| CEN | 0.091 (0.026) | 0.05-0.16 |
| DMN | 0.060 (0.022) | 0.03-0.13 |
| Sub | 0.079 (0.004) | 0.07-0.09 |

Values reflect mean (SD) and range of adjusted R² estimates of structure-function coupling within each resting-state network. “216” and “554” refer to the parcellation resolutions derived from the Schaefer cortical atlas (200- or 500-region versions) combined with the Melbourne subcortical atlas (16 or 54 regions, respectively). Pearson and partial denote the functional connectivity estimation method. CEN = central executive; DAN = dorsal attention; DMN = default mode; Lim = limbic; SMN = sensorimotor; Sub = subcortical; VAN = ventral attention; Vis = visual.

#### Genome-Wide Association Studies (functional annotation and gene mapping):

Functional annotation using FUMA SNP2GENE showed that all lead SNPs across the three regions were noncoding, with modest CADD scores and chromatin states consistent with regulatory function. Gene mapping integrated positional, eQTL, and chromatin-interaction.

Table S10. Lead single-nucleotide polymorphisms in each locus

| SNP | Chromosome:Position (hg build) | Alleles | p | β | SE |
| --- | --- | --- | --- | --- | --- |
| Left precentral gyrus | | | | | |
| rs7914305 | 10:134312761 | C:T | 4.7×10⁻¹⁴ | 0.05 | 0.007 |
| rs149866894 | 8:108606208 | A:G | 1.5×10⁻⁸ | 0.14 | 0.025 |
| rs5792512 | 11:70017491 | A:AC | 2.0×10⁻⁸ | 0.04 | 0.007 |
| rs2863957 | 2:114089551 | A:C | 4.7×10⁻⁸ | 0.05 | 0.008 |
| rs149498207 | 5:71959362 | A:G | 4.9×10⁻⁸ | 0.15 | 0.028 |
| Right frontal pole | | | | | |
| rs7080472 | 10:96012950 | G:T | 9.3×10_-10_ | -0.04 | 0.007 |
| Left temporal pole | | | | | |
| rs41304619 | 10:76743219 | C:T | 3.3×10^-8^ | 0.01 | 0.001 |

Lead SNPs from each genome-wide locus identified in GWAS of regional structure-function coupling associated with bipolar disorder. For each variant, the rsID, genomic position (GRCh37 compatible with FUMA), alleles as defined by FUMA, p-value, effect estimate (β), and standard error (SE) are shown.

Table S11. Functional annotation of lead SNPs (all noncoding)

| SNP | Nearest gene | Distance (bp) | CADD | RegulomeDB | Min chromatin state | Common chromatin state |
| --- | --- | --- | --- | --- | --- | --- |
| Left precentral gyrus | | | | | | |
| rs7914305 | RP11-432J24.5 (intergenic) | 12368 | 1.23 | 5 | 5 | 15 |
| rs149866894 | PGAM1P13 (intergenic) | 53298 | 0.49 | 7 | 9 | 15 |
| rs5792512 | ANO1 (intronic) | 0 | 2.11 | NA | 4 | 15 |
| rs2863957 | PAX8 (intergenic) | 53023 | 3.50 | 7 | 4 | 15 |
| rs149498207 | CTC-347C20.2 (intergenic) | 2865 | 3.64 | 5 | 5 | 15 |
| Right frontal pole | | | | | | |
| rs7080472 | PLCE1 (intronic) | 0 | 4.07 | 6 | 4 | 15 |
| Left temporal pole | | | | | | |
| rs41304619 | KAT6B (intronic) | 0 | 3.01 | 5 | 2 | 5 |

For each lead variant, the nearest gene and its distance, CADD score, RegulomeDB score, and chromatin state metrics (minimum and common states across tissues) are reported.

Left precentral gyrus: Five loci were associated with SFC in the left precentral gyrus. Using MAGMA gene-based analysis, three genes (*FOXF2*, *ANO1*, and *FADD*) were significantly associated after FDR correction (*p_FDR_*<0.05); only *FADD* remained significant following Bonferroni correction. Gene-set and tissue expression MAGMA analyses showed no associations reaching Bonferroni-corrected significance.

FUMA mapped 116 candidate SNPs to 25 genes using positional, eQTL, and chromatin interaction data. *INPP5A* and *PAX8* were mapped by all three methods, *C10orf91* and *ANO1* by positional and chromatin mapping, *FADD* solely by positional mapping, and *PSD4* only via eQTL mapping.

Chromatin interaction mapping identified 429 significant interactions between associated loci and distal genomic regions using brain-specific Hi-C datasets, predominantly representing intra-chromosomal contacts, several localized within adult cortex.

Using GENE2FUNC pipeline in FUMA, expression profiles from GTEx v8 showed that several mapped genes exhibited lower expression in brain-related tissues, including the cortex. Heatmap clustering revealed consistent downregulation across multiple central nervous system regions. No differentially expressed genes were identified across GTEx v8 tissues after Bonferroni correction (*p*<0.05).

Testing for enrichment across multiple curated gene set databases, significant associations were found with 41 unique gene set (*p_FDR_*<0.05), The enriched sets were related to biological processes including immune and cytokine signaling. For example, reactome pathways for interleukin-36 and interleukin-1 signaling, cytokine signaling in the immune system, and multiple GO biological processes related to cytokine-mediated signaling, response to cytokines, and inflammatory responses. Additionally, extracellular matrix and secreted factors, reflected by significant enrichment of multiple NABA categories (Matr isome, Matr isome-associated, Secreted factors).

Right frontal pole: One locus was associated with frontal pole SFC with *rs7080472* as the lead SNP. MAGMA gene-based, gene-set, and tissue expression analyses did not identify any associations reaching Bonferroni-corrected significance. FUMA mapped 8 candidate SNPs to 10 genes, with *PLCE1* and *NOC3L* supported by positional, chromatin interaction, and eQTL mapping. Chromatin interaction mapping revealed 227 significant links between associated loci and distal genomic regions, mainly in stem cells.

Using GENE2FUNC pipeline in FUMA, *PLCE1* was found to be downregulated in brain tissues. No differentially expressed genes were identified across GTEx v8 tissues after Bonferroni correction (*p*<0.05). Enrichment of input genes in gene sets found significant associations with gene sets related to antiviral treatment response, psychiatric medication metabolism, gastrointestinal cancers, drug-induced adverse effects, and CSF Aβ1-42 levels in mild cognitive impairment.

Left temporal pole: One locus was associated with temporal pole SFC, with the *rs41304619* as lead SNP. MAGMA gene-based, gene-set, and tissue expression analyses showed no Bonferroni corrected significance. FUMA identified 3 candidate SNPs mapped to 12 genes. *KAT6B* and *DUPD1* were mapped by FUMA with positional mapping and chromatin mapping but not eQTL mapping. Chromatin interaction mapping revealed 551 significant links between associated loci and distal genomic regions in brain-specific Hi-C datasets. The majority were intra-chromosomal interactions, with several located in adult cortex and other brain regions.

Heatmap revealed under expression of *KAT6B* across multiple brain tissues. Gene-set enrichment analysis identified several significant pathways (*p_FDR_*<0.05), including Wnt and calcium signaling, and multiple protein serine/threonine phosphatase activity-related gene sets. No differentially expressed genes GTEx v8 tissues or enrichment of input genes in gene sets were significantly associated after Bonferroni correction (*p*<0.05).

#### Heritability and Genetic Correlation:

Figure S7. Genetic correlation between structure-function coupling of the precentral gyrus, frontal pole, and supramarginal gyrus, with psychiatric, cognitive, and MRI-derived traits

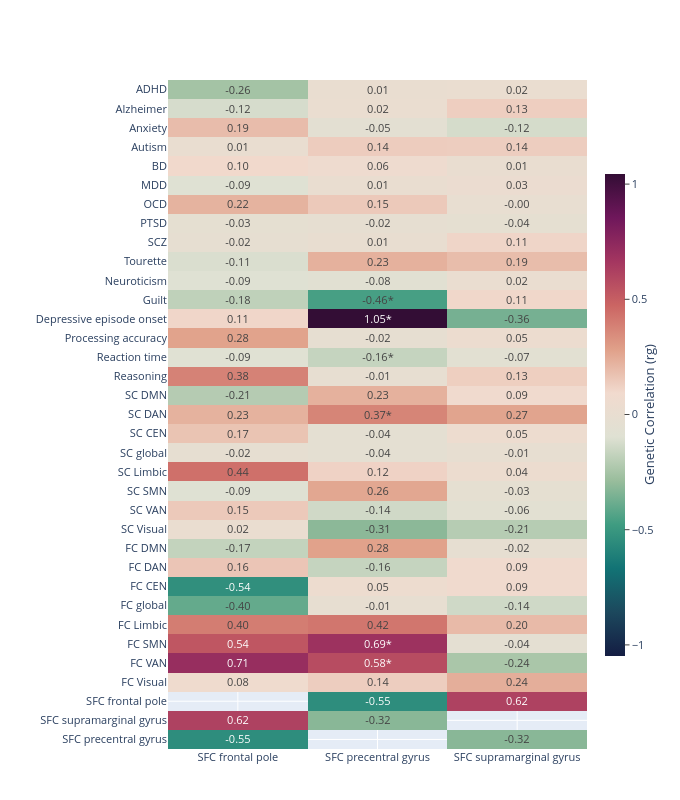

Heatmap displaying LDSC-estimated genetic correlations (rg) between structure-function coupling in the frontal pole, precentral gyrus, and supramarginal gyrus and a set of relevant GWAS traits. Warm colors indicate positive rg values, and cool colors indicate negative rg values. Asterisks (*) denote correlations with *P*<0.05 (uncorrected), reflecting exploratory significance. Values are shown only for SFC traits with sufficient heritable signal. ADHD = Attention-deficit/hyperactivity disorder, BD = Bipolar disorder, CEN = Central Executive Network, DAN = Dorsal Attention Network, DMN = Default Mode Network, FC = Functional connectivity, OCD = Obsessive-compulsive disorder, PTSD = Post-traumatic stress disorder, SC = Structural connectivity, SCZ = Schizophrenia, SMN = Sensorimotor Network, VAN = Ventral Attention Network.
